# Bioinformatics techniques for efficient structure prediction of SARS-CoV-2 protein ORF7a via structure prediction approaches

**DOI:** 10.1101/2022.12.03.518956

**Authors:** Aleeza Kazmi, Muhammad Kazim, Faisal Aslam, Syeda Mahreen-ul-Hassan Kazmi, Abdul Wahab, Rafid Magid Mikhlef, Chandni Khizar, Abeer Kazmi, Nadeem Ullah Wazir, Ram Parsad Mainali

**Affiliations:** Department of Microbiology, Shaheed Benazir Bhutto Women University (SBBWU), Peshawar, Pakistan; Medical Officer, Tehsil Headquarter Hospital, Kallur Kot, Bhakkar, Punjab, Pakistan; Tehsil Headquarter Hospital, Ahmad Pur East, Bahawalpur, Punjab, Pakistan; Shanghai Center for Plant Stress Biology, CAS Center for Excellence in Molecular Plant Sciences, Chinese Academy of Sciences, Shanghai 200032, China; Department of Biotechnology, University of Samarra, Iraq; Institute of Molecular Biology and Biotechnology, University of Lahore, Lahore, Pakistan; Department of Biotechnology, Faculty of Chemical and Life Sciences, Abdul Wali Khan University Mardan (AWKUM), Mardan, Pakistan; Department of Genetics, Institute of Hydrobiology, University of Chinese Academy of Sciences (UCAS), Wuhan, China; Department of Pharmacy, COMSATS University Islamabad, Abbottabad Campus, KPK, Pakistan; National Agriculture Genetic Resource Center (National Genebank), NARC, Khumaltar, Nepal

**Keywords:** protein structure prediction, homology modeling, threading recognition, Ab initio, SARS CoV-2, ORF7a protein

## Abstract

Protein is the building block for all organisms. Protein structure prediction is always a complicated task in the field of proteomics. DNA and protein databases can find the primary sequence of the peptide chain and even similar sequences in different proteins. Mainly, there are two methodologies based on the presence or absence of a template for Protein structure prediction. Template-based structure prediction (threading and homology modeling) and Template-free structure prediction (ab initio). Numerous web-based servers that either use templates or do not can help us forecast the structure of proteins. In this current study, ORF7a, a transmembrane protein of the SARS-coronavirus, is predicted using Phyre2, IntFOLD, and Robetta. The protein sequence is straightforwardly entered into the sequence bar on all three web servers. Their findings provided information on the domain, the region with the disorder, the global and local quality score, the predicted structure, and the estimated error plot. Our study presents the structural details of the SARS-CoV protein ORF7a. This immunomodulatory component binds to immune cells and induces severe inflammatory reactions.

## Introduction

Many organisms’ gene sequences have been determined thanks to considerable efforts in genome sequencing over nearly three decades. Over 100 million nucleotide sequences from over 300,000 different organisms have been deposited in the major DNA databases, DDBJ/EMBL/GenBank, totaling nearly 200 billion nucleotide bases (roughly equivalent to the number of stars in the Milky Way) (Cognato et al., 2020). Over 5 million nucleotide sequences have been converted to amino acid sequences and stored in the UniProtKB database (Chhajed et al., 2023). UniParc has tripled this amount of protein sequences. By the end of 2007, the Protein Data Bank (PDB) included 44,272 protein structures, accounting for less than 1% of the sequences in the UniProtKB database (Stamboulian, 2015).

Proteins are vital molecules that are involved in a variety of biological activities. The majority of protein structure prediction relies on the sequence and structural homology. Protein structure prediction or modeling is critical since a protein’s activity is largely determined by its three-dimensional structure (Lupas et al., 2021). A protein’s 3D structure is also determined by its amino acid content. A slight change in the protein sequence can cause significant structural changes in the native structure (Kryshtafovych et al., 2019). It is critical to have a precise understanding of protein 3D structure, yet interpreting the natural structure of a protein in physiological settings is difficult (Agnihotry et al., 2022). Proteins differ primarily in their amino acid sequence, which leads to differences in spatial shape and structure and biological functions in cells. However, nothing is known about how a one-dimensional sequence of a protein folds into a specific three-dimensional structure. The genetic code uses a triple-nucleotide codon in a nucleic acid sequence to specify a single amino acid in the protein sequence. At the same time, the link between the protein sequence and its steric structure is referred to as the second genetic code (or folding code).

Since the turn of the century, computer approaches for predicting protein structure from its amino acid sequence have sprouted like mushrooms. Anfinsen demonstrated this in 1973 when he published a paper demonstrating that all the information a protein needs to fold properly is encoded in its amino acid sequence, later known as Anfinsen’s dogma (Vila, 2022). The amino acid sequence determines the protein’s fundamental molecular composition from a physical standpoint. Its native structure corresponds to the most stable conformation with the lowest free energy. Although we know that several physical principles regulate the protein folding process, accurately describing such a complex macromolecule is difficult (Deng et al., 2018).

Although it is critical to have a thorough understanding of protein 3D structure, it can be challenging to determine a protein’s natural structure when it exists in the physiological environment of the body. The structure of proteins and protein-ligand complexes is primarily determined using X-ray crystallography and nuclear magnetic resonance spectroscopy (NMR) techniques. These methods for experimentally determining structure are exceedingly time-consuming, expensive, and difficult. As a result, a theoretical understanding of protein structure, dynamics, and folding has been used to construct a model using amino acid sequences with the creation of a computer algorithm and computational tools.

The process by which an individual amino acid sequence folds into a three-dimensional (3D) protein has remained one of the extraordinary unsolved problems for a very long time. Protein sequence and structural data analysis from known and verified structures have investigated the various problems related to amino acid properties, and their arrangement in the form of a helix, sheet, coil, side chain, conformations, and other structural problems related to the packing of sequences into 3D protein models (Pelay□Gimeno et al., 2015, Merritt et al., 2020, Kuhlman and Bradley, 2019, Agnihotry et al., 2022). Numerous novel theories on molecular evolution and methods for structure prediction have been created based on these data and principles. There are several tools for predicting tertiary structures that are based on homology modeling, threading/fold recognition, and the ab initio method (Mathur et al., 2022, Agnihotry et al., 2022).

Engineering unique high-order assemblies, creating protein structures from scratch with new or improved features, and developing signaling proteins with therapeutic potential have all been accomplished using new techniques for constructing protein folds and protein-protein interfaces. A predictive understanding of the relationship between amino acid sequence and protein structure will therefore lead to new opportunities for both the rational engineering of novel protein functions through the design of amino acid sequences with specific structures, as well as for the prediction of function from genome sequence data (Kuhlman and Bradley, 2019).

### 1.0. Protein Structure Predicting Methodologies

The goal of protein structure prediction and engineering design is to close the obvious gap between sequence and structure space. Current protein-predicting methods mainly include template-based and Template-free structure prediction. Template base modeling includes two methods: (i) threading and (ii) homology modeling. At the same time, the template-free prediction approach is also known as ab initio. Protein 3D structures may be easily computed from amino acid sequences using evolutionary information from genomic sequences. These prediction methodologies’ fundamental concepts and advancements will be thoroughly examined (Pandurangan and Blundell, 2020, Ittisoponpisan et al., 2019).

**Figure 1:**
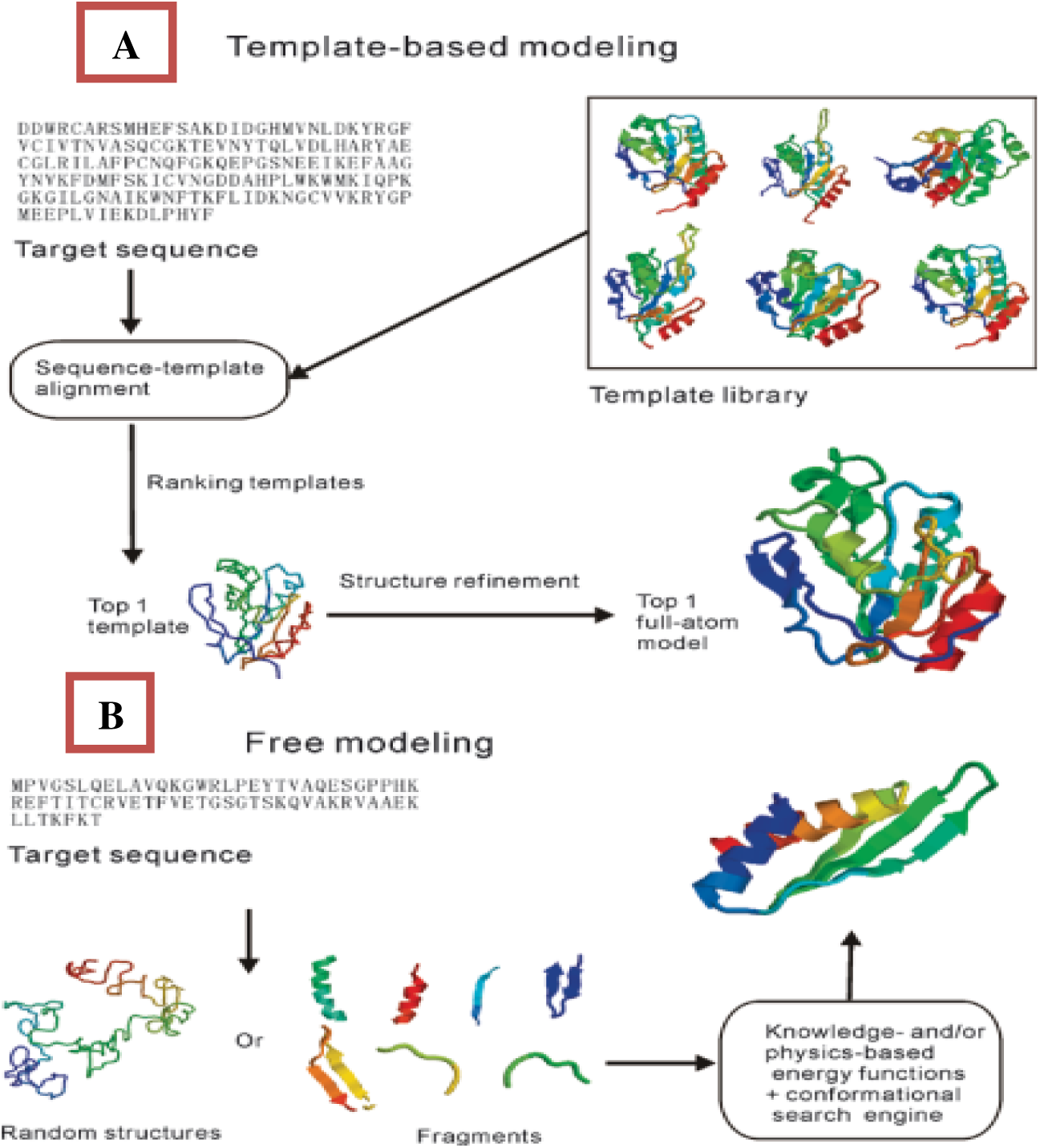
Template-base modeling (a): Sequence of protein targeted template is done by using template library (step 1). Different models for the template are designed (step 2). The top 1 model of the reduced structure is selected which is close to the native structure (step 3). If any changes are required then structure refinement is needed to be done (step 4). The top 1 full atomic model is selected (step 5). Template-free modeling (b): targeted protein is been sequenced first (step 1). Random structures are synthesized and then cut down into fragments (step 2). Based on knowledge-base or Physics-base energy functions and conformational search engine we can conclude the top 1 model (step 3) (Wu and Zhang, 2010)

### 1.1. Template-based Structure Prediction

Protein tertiary structure prediction is commonly made using template-based modeling (TBM), which includes protein threading and homology modeling. TBM uses one or more templates with solved structures to predict the structure of a query protein, referred to as the target protein (Wu and Xu, 2021). Template-based prediction methods construct 3D structures using a template library of solved 3D protein structures (Deng et al., 2018). TBM has always held great promise in terms of producing atomically precise models that are near the native conformation. The target sequence is mapped onto the parent structure first, generating a template structure. Backbone movements, side-chain packing, and loop modeling are all used to develop this hypothetical structure. We would use the following terminology throughout this review for uniformity and clarity. The word “native” refers to the target sequence’s empirically established structure. The protein being modeled is referred to as the “target.” The term “parent” refers to a previously identified protein structure utilized to build the first “template” structure. Finally, any prediction of the native structure is referred to as a “model” structure (Qu et al., 2009, Tunyasuvunakool et al., 2021).

**Figure 2:**
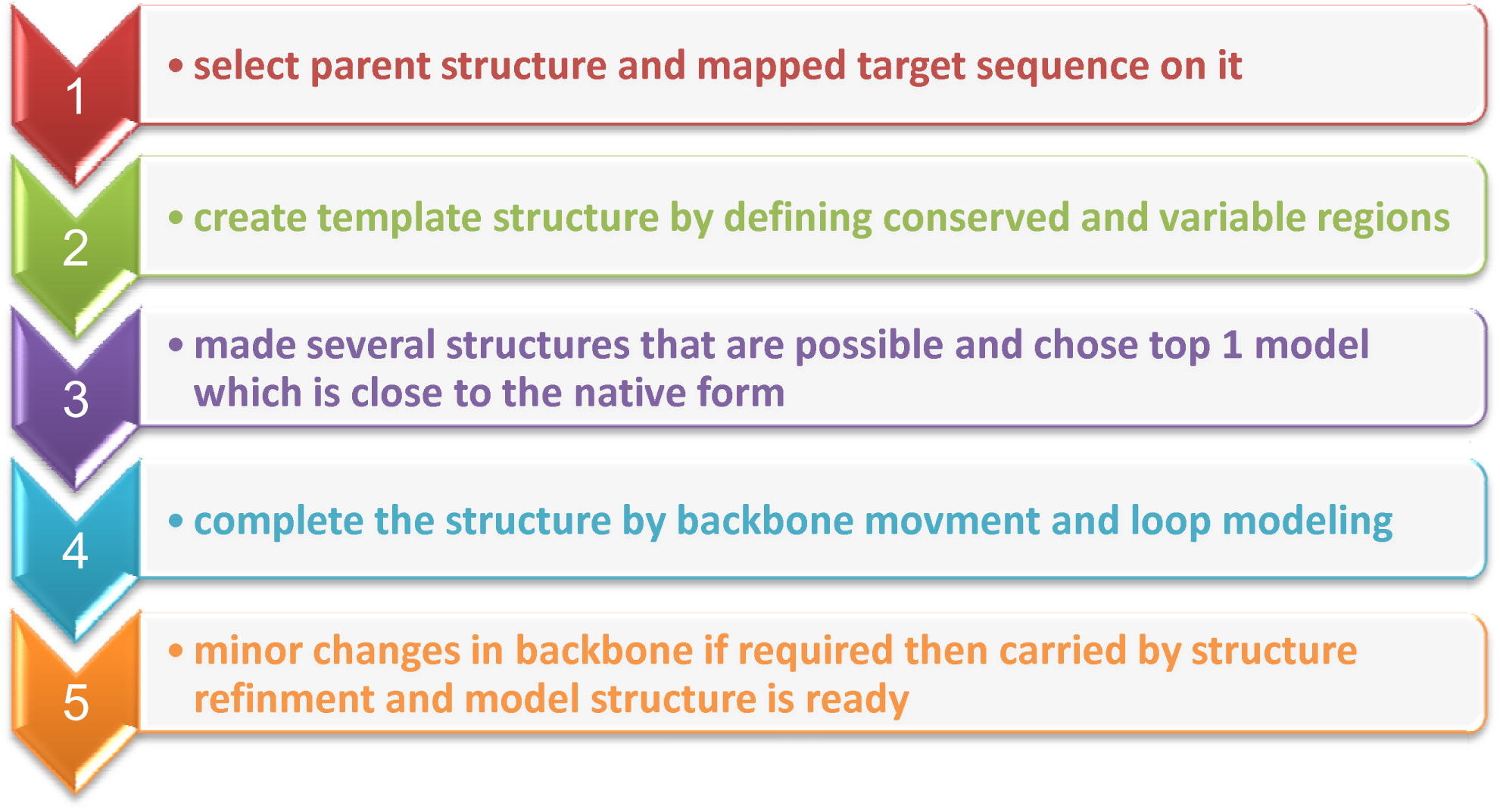
Flow chart for the general procedure of template-base structure prediction

We have arranged our discussion of TBM into the five phases shown in the above flowchart, despite the distinction between steps being arbitrary because many approaches mix steps. TBM approaches include homology modeling based on sequence comparison and threading methods based on fold identification (Zheng et al., 2019).

#### 1.1.1. Homology Modeling

Homology modeling, also known as comparative protein modeling, is the process of creating an atomic-resolution model of a “target” protein using its amino acid sequence and an experimental three-dimensional structure of a similar homologous protein (the “template”). Homology modeling is based on finding one or more known protein structures that are likely to mimic the structure of the query sequence and creating an alignment that maps query sequence residues to template residues (Yadav and Mohite, 2020).

The basis of homology modeling is the finding that identical sequences from the same evolutionary family generally adopt similar protein structures. It is currently the most accurate method of predicting protein structure using a homologous structure in the PDB as a template. With the continuous expansion of the PDB database, homology modeling can now predict an increasing number of target proteins (Wang et al., 2008, Deng et al., 2018).

Template selection, target-template alignment, model development, and model assessment are the four processes in the homology modeling process. Because the most prevalent techniques of finding templates rely on creating sequence alignments, the first two phases are frequently combined. The information included in a template and alignment must be employed to construct a three-dimensional structural model of the target, expressed as a collection of Cartesian coordinates for each atom in the protein.

##### General steps for Homology modeling

Following are the general steps for homology modeling:

1. **Template selection:** Determining the appropriate template structure is a critical initial step in homology modeling. Sequence alignments, supplemented by database search tools like FASTA and BLAST, are the most basic way of template identification.
2. **Target-template alignment:** Upload the main sequence to fold-recognition servers, which will recognize similarities and align the target sequence according to the template. When running a BLAST search, a good initial step is to look for matches with a low E-value that are considered close enough in evolution to create a reliable homology model.
3. **Model development:** The information included in a template and an alignment must be utilized to construct a three-dimensional structural model of the target, represented as a set of Cartesian coordinates for each atom in the protein.
4. **Model assessment:** Statistical potentials or physics-based energy estimates are the most common approaches for evaluating homology models without reference to the target structure. Both approaches yield an energy estimate for the model or models being evaluated (Hasani and Barakat, 2017).

**Figure 3:**
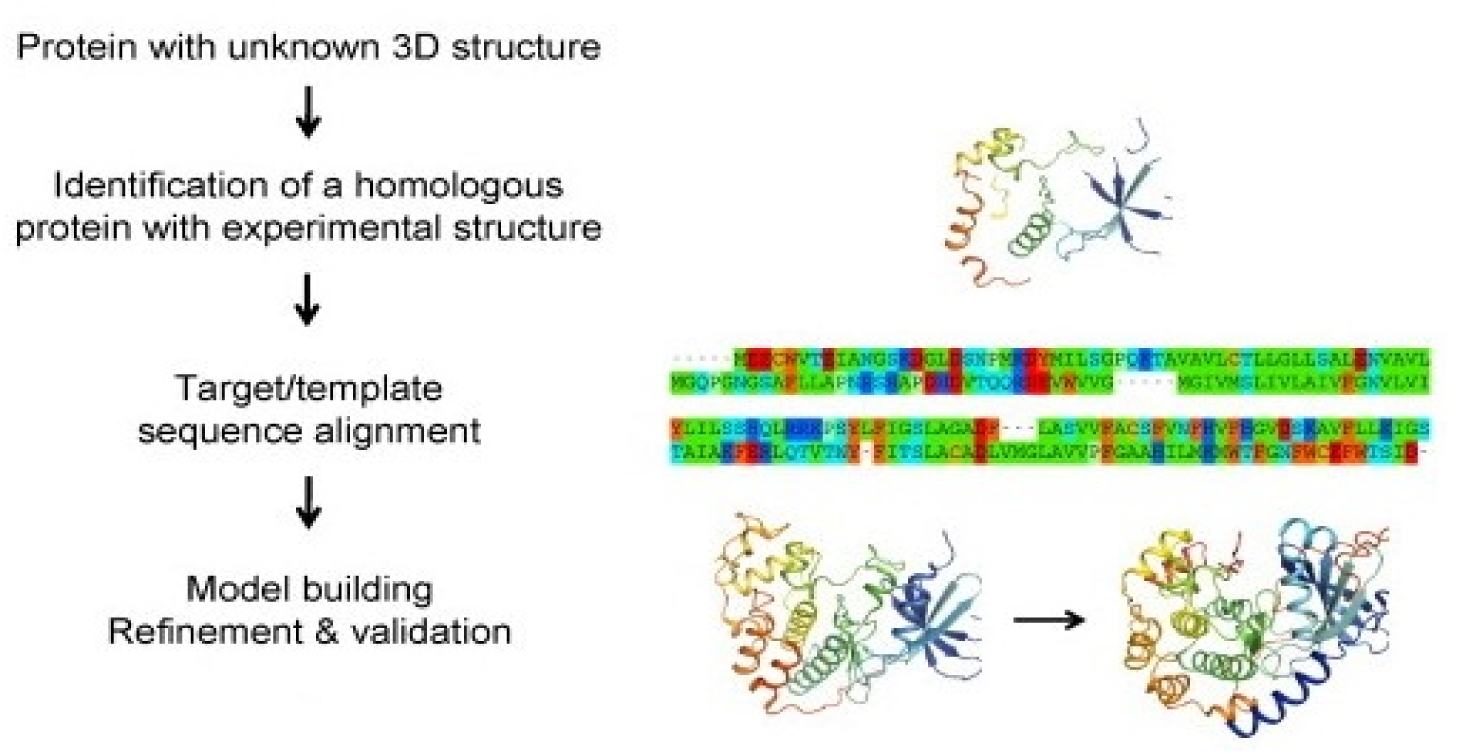
Basic steps of the Homology technique (Cavasotto and Phatak, 2009)

**Table 1:** Software used in Homology modeling (List of protein structure prediction software, March 2022)

| Description | Method | Websites |
| --- | --- | --- |
| SWISS-MODEL | Local similarity/fragment assembly | https://swissmodel.expasy.org/ |
| Yasara | Detection of templates, alignment, and modeling including. ligands and oligomers, hybridization of model fragments | http://www.yasara.org/ |
| MODELLER | Satisfaction of spatial restraints | https://salilab.org/modeller/ |
| MOE (Molecular Operating Environment) | Template identification, use of multiple templates, loop modeling, rotamer libraries for side-chain conformations, relaxation using MM forcefields | https://www.chemcomp.com/Products.htm |

#### 1.1.2. Threading recognition

Protein threading, also known as fold recognition, is a protein modeling technique used to model proteins that have the same fold as proteins with known structures but are not homologous (Zheng et al., 2021). Threading involves matching the target sequence to homologous and distant-homologous structures using an algorithm and selecting the best matches as structural templates. The underlying concept of threading is that protein structure has evolved to be very conservative, and the number of distinct structural folds in nature is restricted. When no structures with evident sequence similarity to the target protein are located in PDB, proteins having structural similarity to the target protein can still be detected (Schauperl and Denny, 2022).

Threading is based on two principles: (i) identical basic sequences fold into similar structures, and the number of possible structural folds of proteins is finite, meaning that even non-homologous proteins can have similar structures. In this method, the target protein’s amino acid sequence is “threaded” onto the structure of a suitable template, followed by local structural rearrangement and refinement (Nikolaev et al., 2018).

The prediction is made by “threading” (i.e., aligning) each amino acid in the target sequence to a place in the template structure and evaluating how well the target fits the template (Källberg et al., 2012). Following the selection of the best-fit template, the structural model of the sequence is constructed using alignment with the chosen template (Källberg et al., 2012, Zhou et al., 2022).

##### The general procedure of the threading technique

The following are the four steps in a generic protein threading paradigm:

1. **Sequence cutting and threading:** The target sequence has been cut down into segments and then threaded to form structural domains.
2. **Construction of a structure template:** Select protein structures as structural templates from protein structure databases. Protein structures are often chosen from PDB, FSSP, SCOP, or CATH databases.
3. **Scoring function design:** Create a scoring function (A scoring formula is used to determine the chance that a model structure is a native structure) to determine the fitness of target sequences and templates based on known correlations between structures and sequences. The accuracy of prediction, particularly alignment accuracy, is highly connected to the quality of the energy function.
4. **Threading alignment:** Using the provided scoring function, align the target sequence with each structural template. This phase is one of the most important tasks of any threading-based structure prediction system.
5. **Threading prediction:** As the threading prediction, choose the threading alignment that is statistically most likely. Then, the target sequence’s backbone atoms and the selected structural template’s aligned backbone locations create a structure model for the target (Batool et al., 2019).

**Figure 4:**
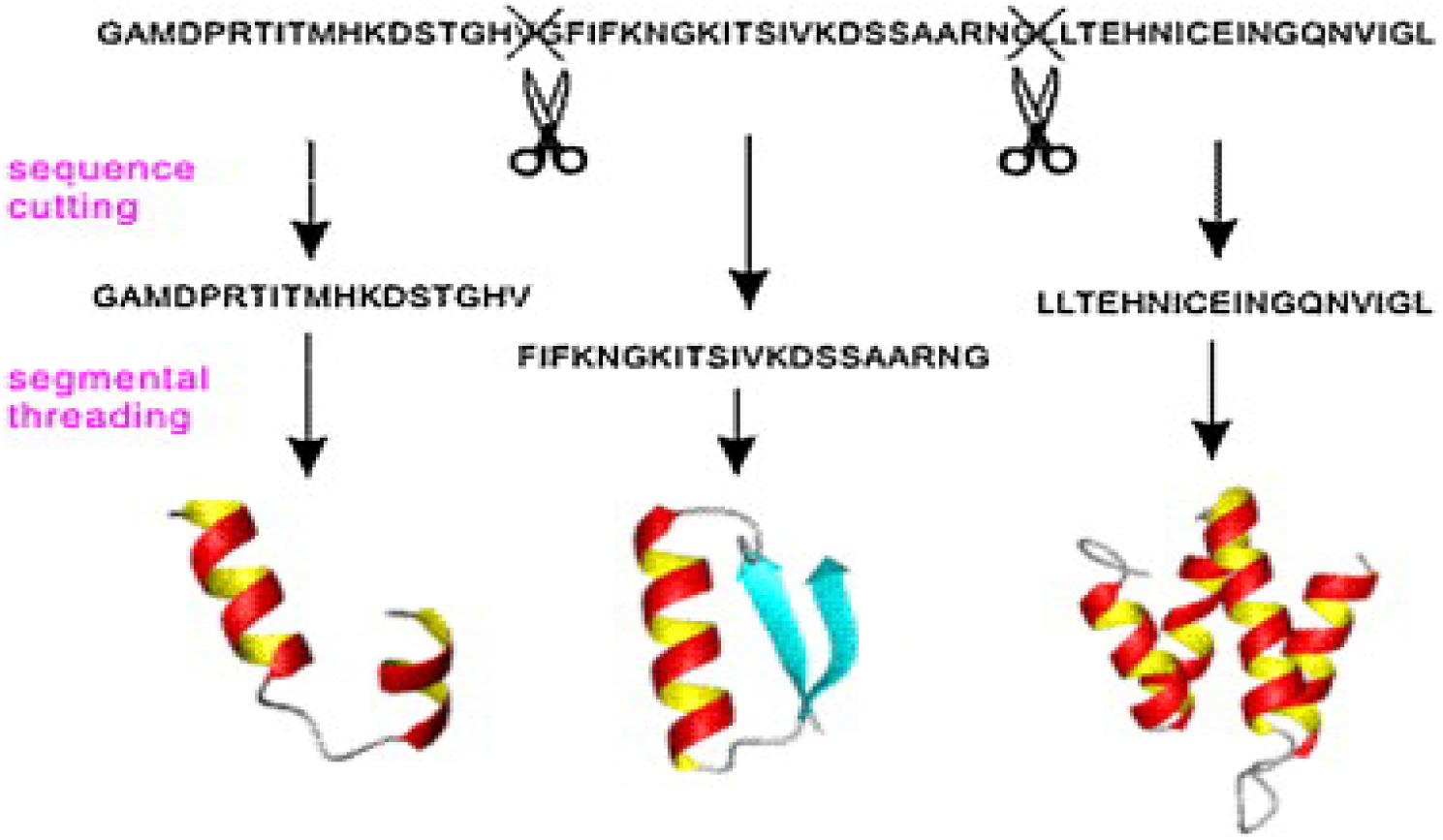
Cutting of target sequence into fragments and each fragment constitute a secondary structure (Wu and Zhang, 2010)

**Figure 5:**
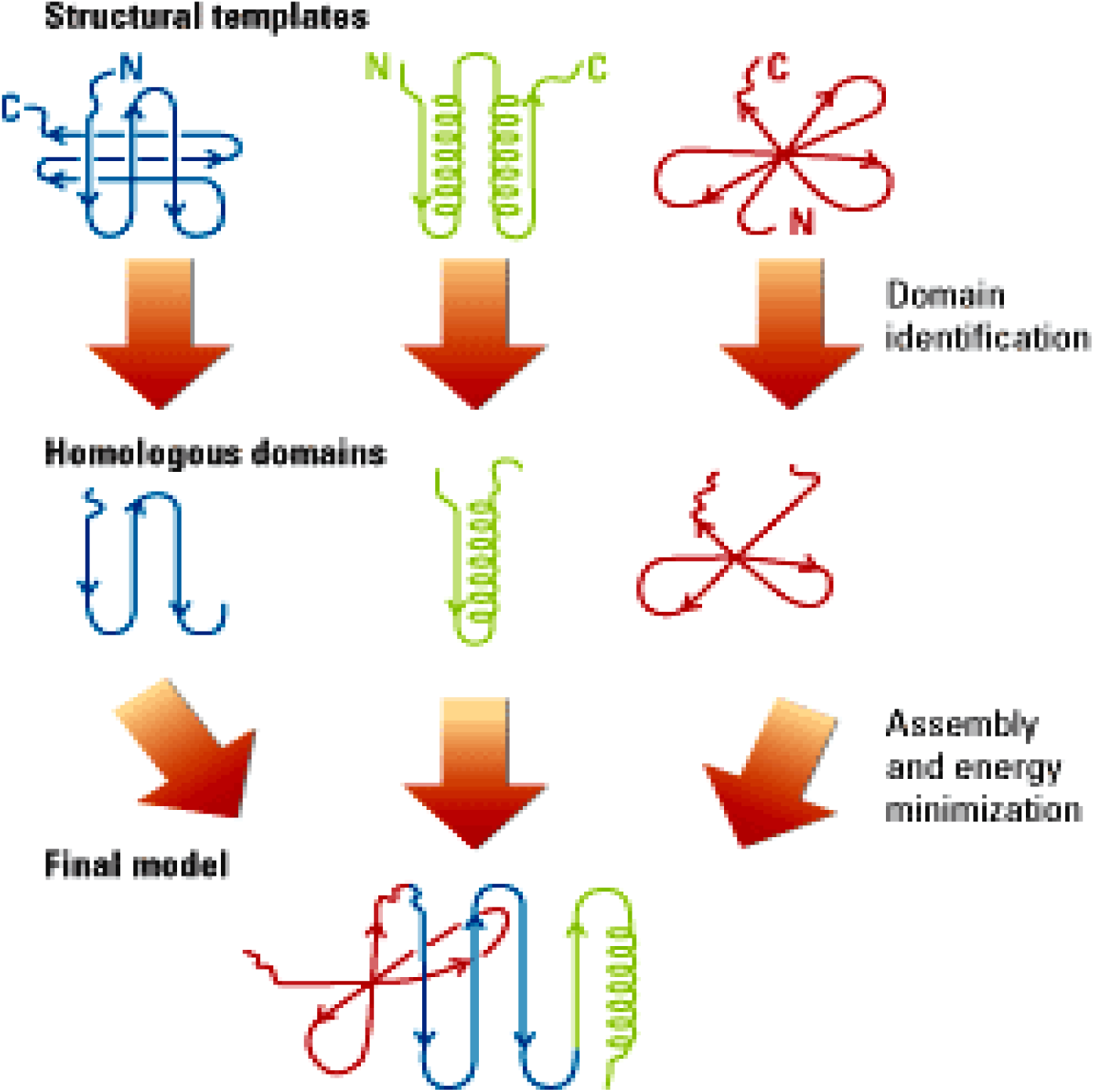
Structural templates and homologous domains are merged to form the final model (Holliman, 2000)

**Table 2:**
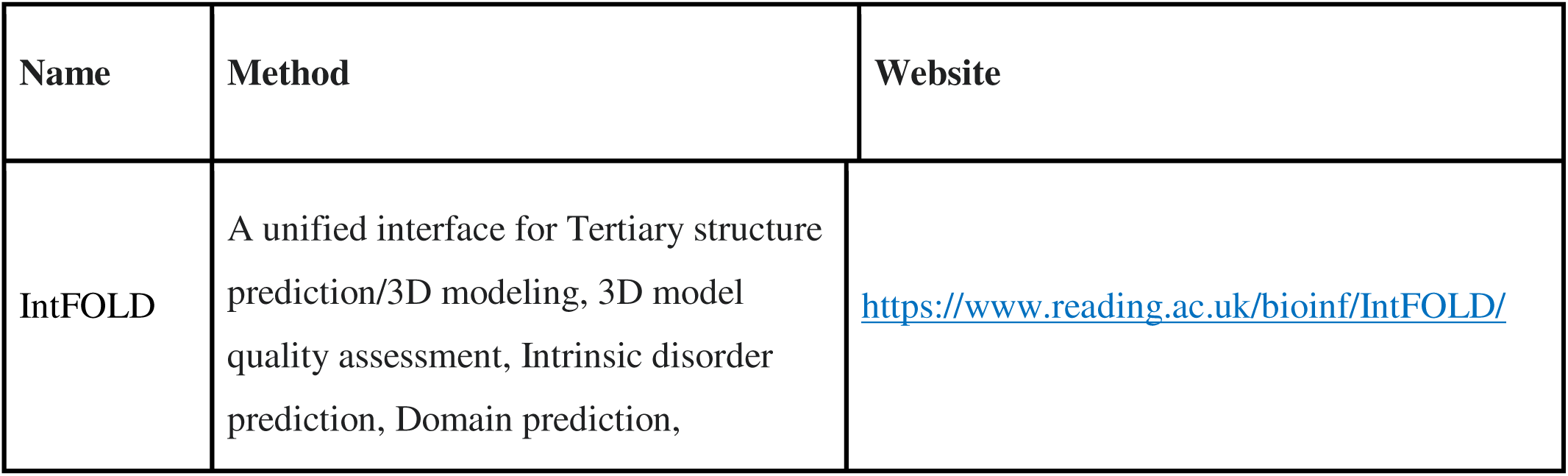
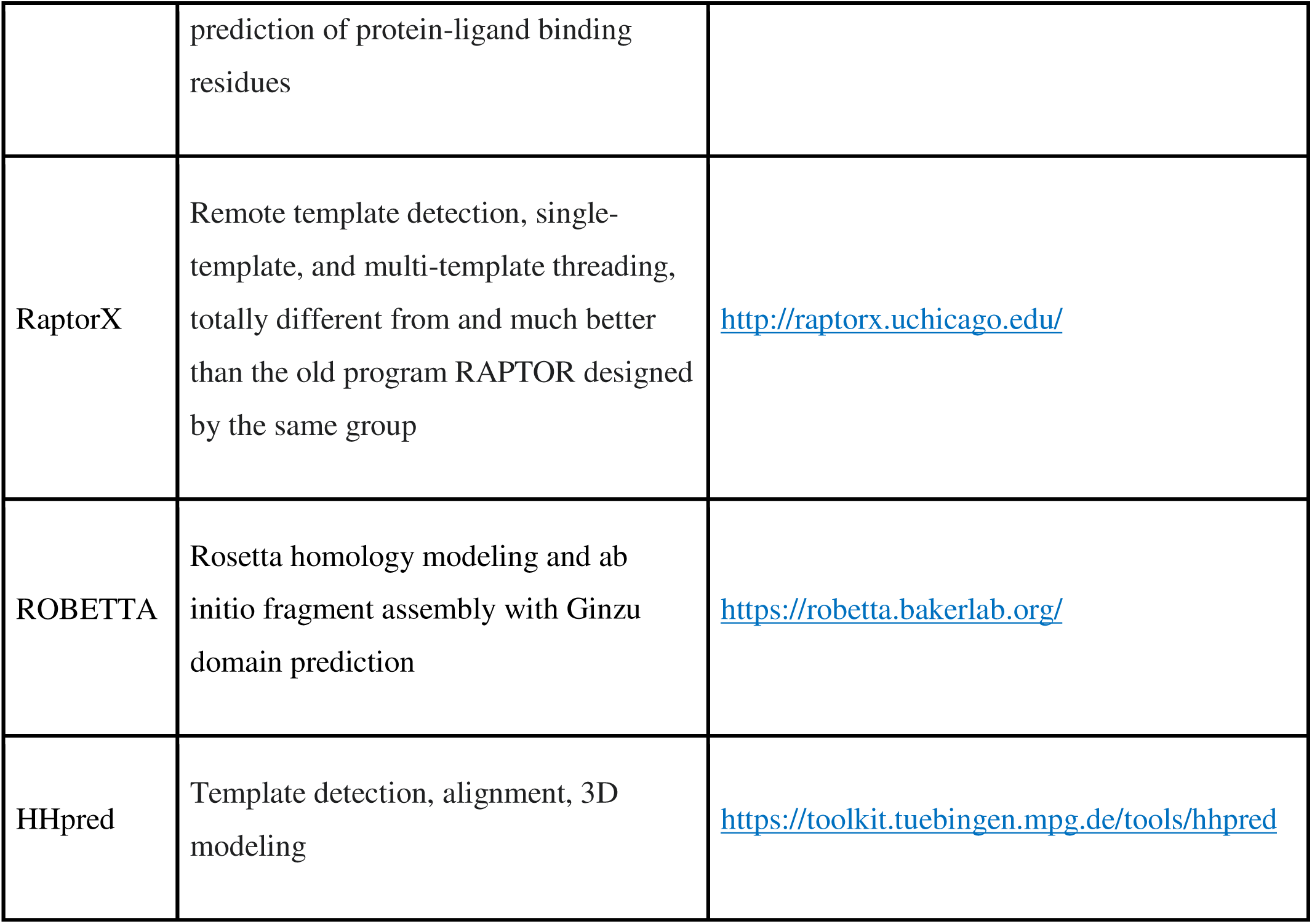
Software use in threading modeling (List of protein structure prediction software, March 2022)

### 1.2. Template-free Structure Prediction

When structural analogs are not found in the PDB library, or threading fails to identify them, the structure prediction must be created from scratch. This prediction is known as ab initio or de novo modeling, which roughly translates to “modeling from the ground up.” Although free modeling is the most efficient method, hybrid models that include both knowledge-based and physics-based potentials should be considered (Dhingra et al., 2020). Unlike homology modeling and threading, the ab initio technique tries constructing structures from the base up without relying on any already solved structures. Currently, to some extent, nearly all protein structure prediction algorithms employ structural information from already solved structures. As a result, the term “template-free” is frequently used to describe approaches that are neither homology modeling nor threading (Dhingra et al., 2020).

Ab initio modeling usually involves a conformational search guided by a designed energy function (the total energy of a system is expressed as a function by the energy functional). Typically, this technique creates many alternative conformations (also known as structural decoys), from which final models are chosen. As a result, three criteria are required for successful ab initio modeling: (i) an accurate energy function, using which the native structure of a protein corresponds to the most thermodynamically stable state among all possible decoy structures; (ii) an efficient search method, using conformational search, to quickly identify low-energy states; a strategy for selecting near-native models from a pool of decoy structures (Lei et al., 2021, Lee et al., 2017).

#### The general procedure of the Ab initio method

Following are the general steps for Ab initio/template free prediction method:

- **Query sequence:** query sequence refers to the target sequence needed to predict.
- **Fragment library:** query sequence cut down in the fragments and synthesizes secondary structure in the form of pieces. Thus more than one fragment is formed, so the whole set is known as a fragment library.
- **Fragment assembly:** fragments are assembled by different methods that work with the help of energy function. Some of the best-known methods include Fragfold, Simfold, and Rosetta.
- **Low and high-resolution models:** low-resolution models are first formed with backbone Cα-atom. In contrast, after all-atom reconstruction, high-resolution models are generated.
- **Final model:** After the high-resolution model, we have to select one close to the native conformation by the Monte Carlo approach or Critical assessment of structure protein (CASP).

**Figure 6:**
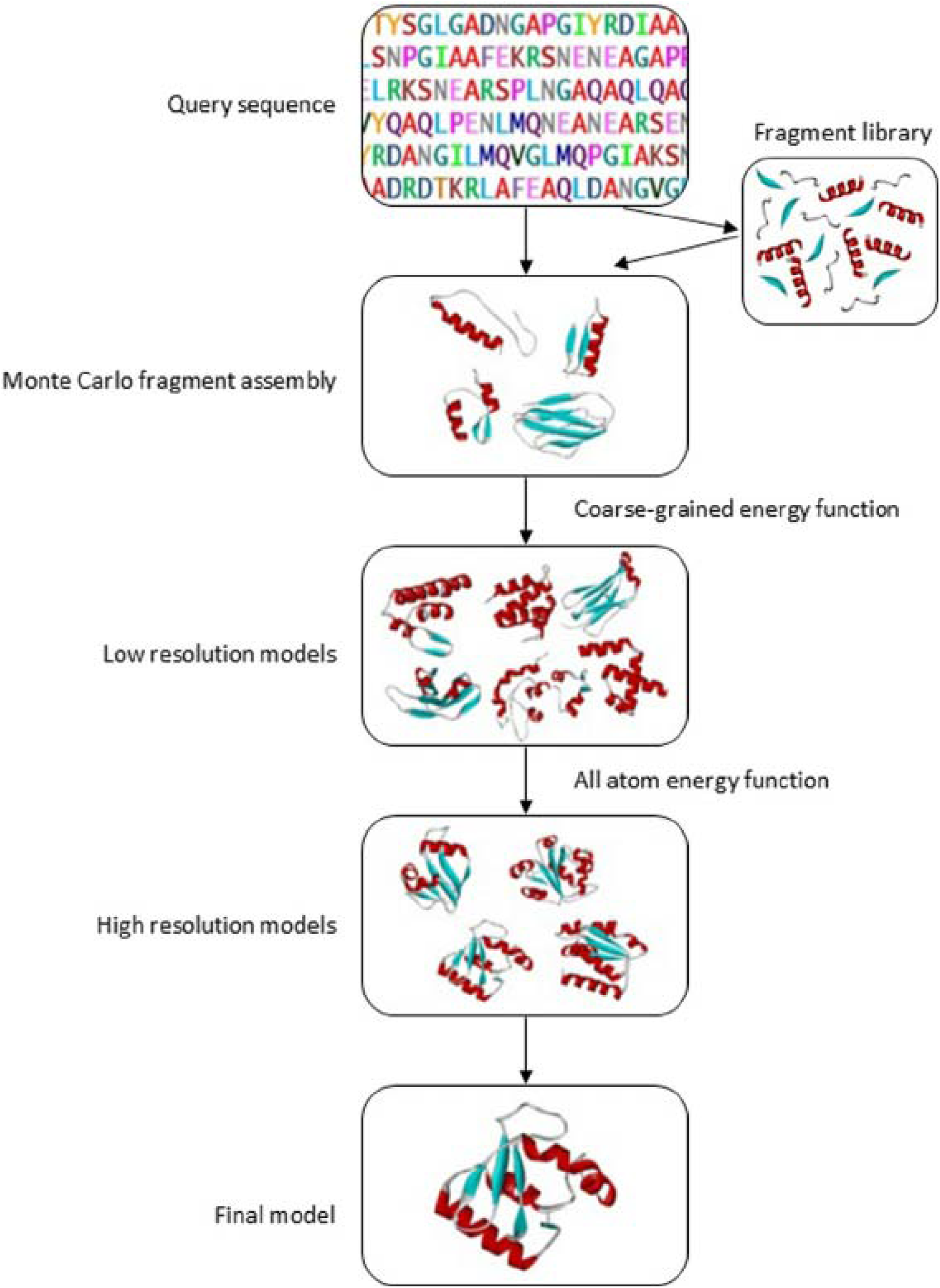
Basic procedure followed in Ab initio approach (Khor et al., 2015)

**Table 3:** Software used in Ab initio (List of protein structure prediction software, March 2022)

| Name | Method | Website |
| --- | --- | --- |
| trRosetta | It builds the protein structure based on Rosetta. It also includes inter-residue distance and orientation distributions, predicted by a deep residual neural network. | https://yanglab.nankai.edu.cn/trRosetta/ |
| I-TASSER | Threading fragment structure reassembly | https://zhanggroup.org/I-TASSER/ |
| ROBETTA | Rosetta homology modeling and ab initio fragment assembly | https://robetta.bakerlab.org/ |
| Abalone | Molecular Dynamics folding | http://www.biomolecularmodeling.com/Abalone |

**Figure 7:**
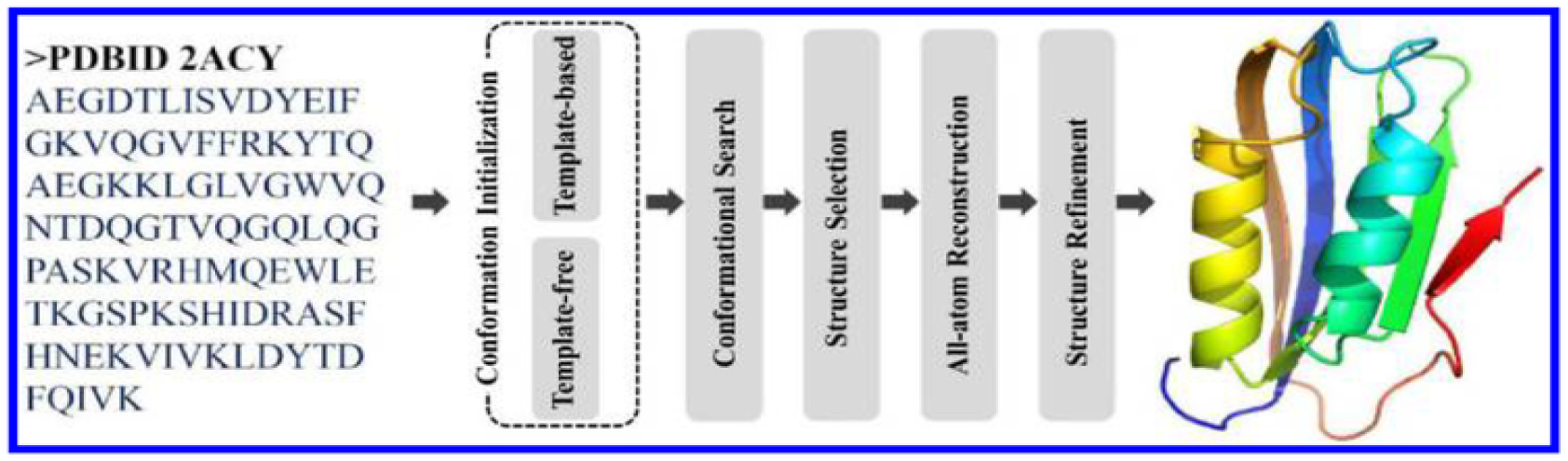
Mechanism of protein structure prediction (Deng et al., 2018)

### 2.0. General Steps of Protein Structure Prediction

Although different prediction approaches use detailed prediction procedures, the core phases are always the same, including conformation initialization, conformational search, structure selection, all-atom structure reconstruction, and structure refining. These processes are described, along with some well-known approaches or instruments that are widely employed in each. We will also briefly explain the current challenges and future directions in protein structure prediction.

#### 1. Conformation initialization

The one-dimensional amino acid sequence of the target protein serves as the beginning point (input) for protein structure prediction. The model of three-dimensional structures serves as the final output. Although a protein sequence is theoretically potential steric conformation is nearly limitless, the native one for most proteins is unique (Deng et al., 2018, Yang et al., 2018).

The manner of conformation initialization is the main distinction between the “template-based” and “template-free” systems. The template-based technique finds the first conformation by looking for homologous or structurally comparable solved structures to the target protein. The first conformation is generally built via fragment assembly in the template-free technique (Kuhlman and Bradley, 2019).

Generally, the initial conformation obtained from a structural template is vastly better than anyone built from scratch. It can dramatically shorten the process of subsequent conformational search. However, there is no guarantee that satisfactory structural templates for any target protein can always be found (Deng et al., 2018).

Template-free techniques are the best option for challenging target proteins for which no appropriate template can be found. It is the simplest technique to produce the starting conformation of the target protein at random. First, estimate the target protein’s structural properties, such as backbone dihedral angle, solvent accessibility, contact map, or secondary structure, independent of any structural template. Several approaches do not require a template.

Fragment assembly is a method for predicting protein structure, in which matching structural fragments are cut from experimental structures based on secondary structure, backbone dihedral angle, and other factors (Deng et al., 2018, Senior et al., 2020).

#### 2. Conformational search

After constructing the initial conformation, we may conduct calculations with the help of a force field (a computational method used to measure forces among atoms within molecules and between molecules) to look for near-native conformations one by one. A simplified representation of protein conformations is especially important for speeding up protein folding simulation. The structural template identified by sequence alignment already has a reduced conformation with only the backbone or Cα-atoms (Wang et al., 2008). Almost all protein structure assembly simulation techniques currently use a simplified model to perform a conformational search. Each residue, for example, can be represented only by its Cα-atom and the virtual center of the side chain, or a sequence of dihedral angles can represent the complete backbone conformation (Webb and Sali, 2016, Alapati et al., 2020).

Conformational search requires a force field depicting the protein conformational energy landscape. The energy landscape corresponds to any configuration of the target protein. The natural shape of the target protein should be at the lowest point of the energy landscape in a well-designed force field. Many existing force fields are simply based on classical physics theory, which includes bond-stretching energy, angle-bending energy, angle torsional energy, and other non-bonded interaction energies. Several force fields are commonly used, for example, AMBER and CHARMM. A physics-based energy function is another name for them (Guterres et al., 2022, Fan et al., 2021).

On the other hand, the alternative energy function is generated entirely from structural feature information acquired from experimentally solved structures, known as the knowledge-based energy function. It is a dependable structural information pool for building knowledge-based energy functions and a structural template library for homology modeling. These two types of energy functions each have their own set of benefits and drawbacks (Banerjee et al., 2020).

Although physics-based energy functions are built on the fundamental principle and have clear physical importance, they are often inaccurate in characterizing complex atomic interactions and perform poorly in protein structure prediction. Knowledge-based energy functions extensively use experimentally solved structures and provides promising results in many circumstances. However, they are “black boxes” that cannot assist us in understanding the nature of the protein folding process. In reality, most structure prediction algorithms use physics-based and knowledge-based energy functions (Deng et al., 2018).

After determining the energy function, we must devise a method for locating the target protein’s lowest-energy conformation. Conformational search is a type of molecular dynamics simulation that predicts protein folding and movement by solving Newtonian motion equations (F=ma). However, the need for computational resources for molecular dynamics modeling is excessive for biomacromolecules such as proteins. The conformational search approach based on Monte Carlo simulation, which changes the conformation randomly depending on multiple motions planned, can be much quicker. Shifts and rotations of structural segments and position changes of a single atom are examples of these motions. They might be continuous in space or restricted to a cubic lattice structure (Afrasiabi et al., 2022).

#### 3. Structure selection

A huge number of target protein structures are created when the conformational search is completed. One of the unresolved problems in molecular dynamics and Monte Carlo simulation is that conformations are usually stuck at the local minimum state (the first structure formed when energy is emitted) (Kantarci-Carsibasi et al., 2008, Xu and Zhang, 2012). Because of the shortcomings of the force field, even when the minimum global state (most stable conformational structure of the protein) is found, the conformation does not always match the one that is closest to the native state.

The assessment method for identifying native-like structures from non-native structures is a critical aspect of structure selection. CASP (Critical assessment of structure prediction) has a unique prediction category for evaluating structural quality assessment methodologies. Various structural decoys, including CASP structural models, are available for training and testing methods of structural quality assessment (Moult et al., 2014, Kryshtafovych et al., 2019, Kryshtafovych et al., 2021).

In essence, the force field can filter structures, and the output of conformational search is simply the force field-filtered structures. We may use a lot more sophisticated energy functions for structure selection. The energy function can be based on physics or knowledge, although the latter is more common and successful.

#### 4. All-atom structure reconstruction

Most prediction approaches use simplified protein representations for conformational search; thus, models should be used to recreate the all-atom structure. For reduced models based on different protein representations, the method of all-atom reconstruction differs greatly. Some prediction algorithms use the “Cα atom” plus “virtual center of side chain” representation, where the “virtual center of side chain” serves as an aid in establishing the position of the Cα atom during the conformational search (Rotkiewicz and Skolnick, 2008, Pan et al., 2021, Deng et al., 2018).

The reconstruction procedure is frequently split into two halves in this situation. The first step is to recreate the backbone atoms (C N and O) using the positions of Cα atoms, which is the basic purpose of many all-atom reconstruction approaches. REMO, for example, created a backbone isomer library of 528798 pieces with four consecutive residues collected from 2561 protein chains in PDB. The second stage is to reconstruct each residue’s side chain. The optimal technique to rebuild the side chain is based on the same library as the strategy used to rebuild backbone atoms. Scwrl, SCATD, RASP, and other approaches for side chain reconstruction are available (Badaczewska-Dawid et al., 2020).

#### 5. Structure refinement

Although the previous procedures generated the full structure of the target protein, the structural quality is frequently poor due to flaws in the force field, conformational search, or all-atom reconstruction. Some methods combine the procedures of all-atom reconstruction and refinement, refining the reduced model (such as backbone structure) and all-atom structure separately. According to the reconstruction schedule, structural issues in the reduced model can directly affect the quality of the final all-atom structure (Badaczewska-Dawid et al., 2020).

Structure refinement also requires the use of a force field to run molecular dynamics simulations or Monte Carlo simulations. In structure assembly simulations, conformational search aims to find the target protein’s backbone structure. The major goal of structure refinement is to increase the quality of all-atom structures (particularly local structures) where only minor changes in backbone conformation are made.

Structure refinement involves far fewer conformational changes than structure construction simulations. FG-MD is one of these atomic-level molecular dynamics simulation-based structure refinement approaches. Mod-Refiner is another Monte Carlo simulation-based method for structural reconstruction and refinement (Deng et al., 2018).

### 4.0 Structure Prediction of ORF7a protein of SARS coronavirus

The severe acute respiratory syndrome (SARS) coronavirus, often known as SARS-CoV, was the cause of the worldwide pandemic that resulted in almost 800 fatalities in 2003 (Shereen and Kazmi, 2020, Shereen et al., 2020b, Rafiq et al., 2021). A worldwide outbreak of SARS was traced to the SARS coronavirus in 2003. As of 23 October 2020, the severe acute respiratory syndrome coronavirus 2 (SARS-CoV-2)-caused pandemic COVID-19 epidemic had been linked to 41,570,883 confirmed cases and 1,134,940 fatalities worldwide. Understanding the crucial roles of SARS-CoV macromolecules has been made possible by the availability of their structures. Four structural proteins, sixteen non-structural proteins, and eight accessory proteins are all encoded by its vast RNA genome (Shereen et al., 2020a).

Monocytes are innate immune cells participating in phagocytosis, antigen presentation, and cytokine release during or after migration into tissues and lymphoid organs. They are also involved in other complicated functional processes of inflammatory reactions. An observational investigation of 54 SARS-CoV-2-infected patients revealed that all of these patients had signs of severe immunological dysregulation caused by dysfunctional monocytes. Proteins with immunoglobulin-like (Ig-like) domains are critical mediators of immune system macromolecular interactions. According to prior research, the SARS-CoV-2 Ig-like viral protein ORF7a interacts with immune cells through its association with integrin LFA-1 (lymphocyte function-associated antigen-1 is a key T cell integrin that plays a major role in regulating T cell activation and migration). It is anticipated that SARS-CoV-2 ORF7a, like SARS-CoV ORF7a, will belong to the Ig-like domain superfamily (Shereen et al., 2020a, Shereen et al., 2021, Zhou et al., 2021).

The ORF7a gene produces an accessory protein, a type I transmembrane protein with 122 amino acids, a 15-residue N-terminal signal peptide, an 81-residue luminal domain, a 21-residue transmembrane region, a 5-residue cytoplasmic tail, according to sequence analysis predictions. Although the ORF7a sequence has been found in all SARS-CoV isolates isolated from human and animal sources, it seems exclusive to SARS. It resembles other viral or non-viral proteins (Elaswad et al., 2020, Addetia et al., 2020, Neches et al., 2021).

SARS-CoV-2 ORF7a’s immunoglobulin-like fold ectodomain interacts with CD14+ monocytes in human peripheral blood with high efficiency. It is significant to note that SARS-CoV-2 ORF7a coincubation with CD14+ monocytes ex vivo resulted in a reduction in HLA-DR/DP/DQ expression levels and an upregulation of proinflammatory cytokine production, specifically IL-6, IL-1, IL-8, and TNF-α. SARS-CoV-2 ORF7a is an immunomodulating factor that binds to immune cells and induces severe inflammatory reactions (Liu et al., 2022, Zhou et al., 2021).

**Figure 8:**
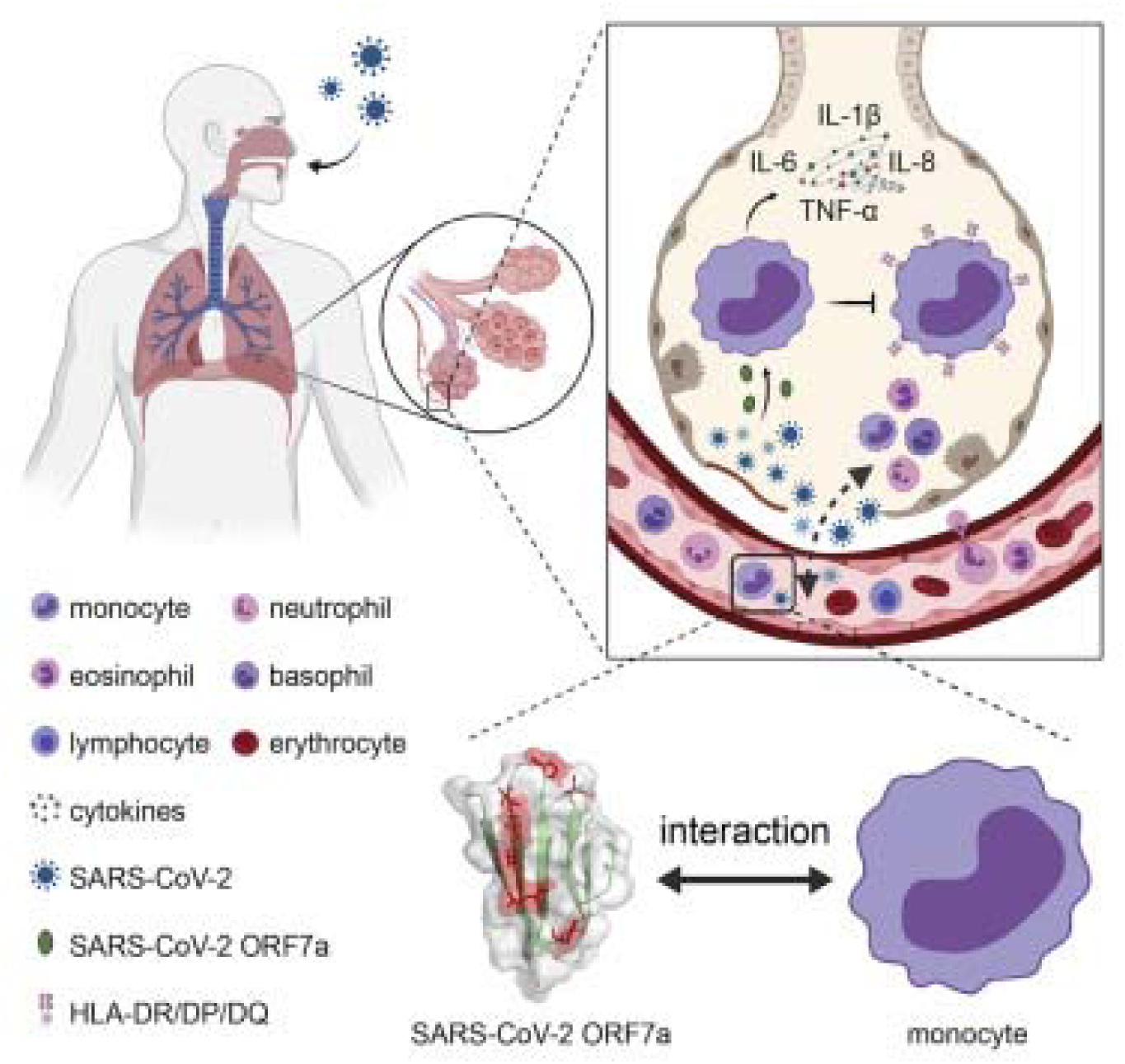
Interaction between monocytes and SARS-CoV-2 ORF7a (Zhou et al., 2021)

There are several protein prediction softwares and web-based services through which we can find the ORF7a protein sequence and predict its protein structure. Some software is based on homology modeling, threading technique, and ab initio simulation. Some software and their results are mentioned below based on these three methodologies.

### 4.1 Phyre & Phyre2

Free web-based resources for protein structure prediction include Phyre and Phyre2, based on Protein Homology modeling and pronounced “fire” (Shelake et al., 2016). Dr. Lawrence Kelley created and wrote the Phyre2 web server (Bennett-Lovsey et al., 2008). Phyre has been mentioned more than 1500 times, making it one of the most widely used techniques for predicting protein structures. It can frequently provide trustworthy protein models, much like other distant homology identification methods (Bakhtin et al., 2019, Kelley, 2017).

The Phyre and Phyre2 services use homology modeling concepts and methods to forecast a protein sequence’s three-dimensional structure. A protein sequence of interest (the target) can be modeled reasonably accurately on a very distantly related sequence of a known structure (the template), provided that the relationship between target and template can be determined through sequence alignment, as the structure of a protein is more conserved in evolution than its amino acid sequence. The initial Phyre server, made available in June 2005, uses a profile-profile alignment technique based on a scoring matrix unique to each protein’s location. The Phyre2 server, which replaced the original Phyre server and offered greater capabilities and a more sophisticated User experience, was made available to the public in February 2011 (Kelley et al., 2015, Jameel et al., 2022).

#### 4.1.1 Methodologies

All species’ genomes, proteins, and sequences are available on the NCBI database. A 122 amino acid ORF7a sequence, an accessory protein of the SARS coronavirus, is acquired from NCBI and entered into the Phyre2 search field. Alternative names for ORF7a protein accessory protein 7a and X4 protein. Results will be dispatched after waiting approximately 30-40 minutes on the website, and as well as it will be emailed too.

#### 4.1.2 Results

The single highest-scoring template has modeled 84 residues (69% of your sequence) with 100% confidence. The model is created by using the c1yo4A template. The c1yo4A template is used for ORF7a and displays 100% alignment coverage.

**Figure 9:**
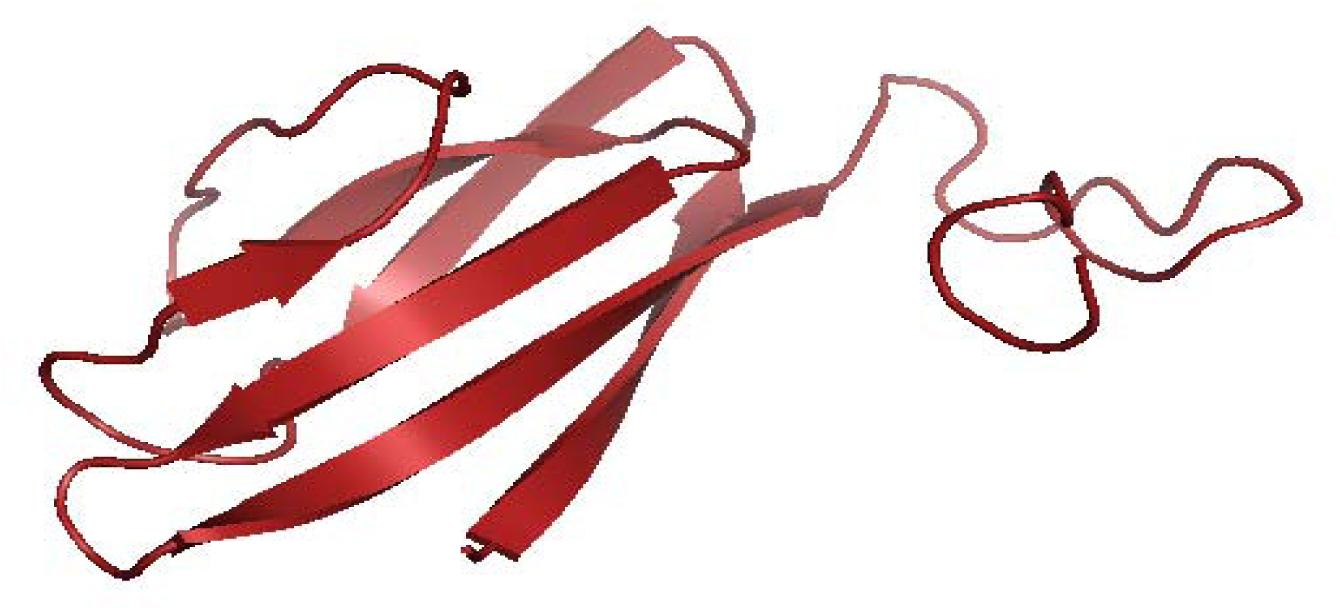
phyre2 results for ORF7a protein

Phyre2 gives us a full model in 3 dimensions along with sequence analysis, sequence analysis, secondary structure and disorder prediction, domain analysis, and template alignment details. Nevertheless, due to only 69% coverage, we cannot consider this model the best-predicted model.Phyre2 detects sequence homologs with PSI-Blast. The ability to accurately and confidently detect homology is common.

**Figure 10:**
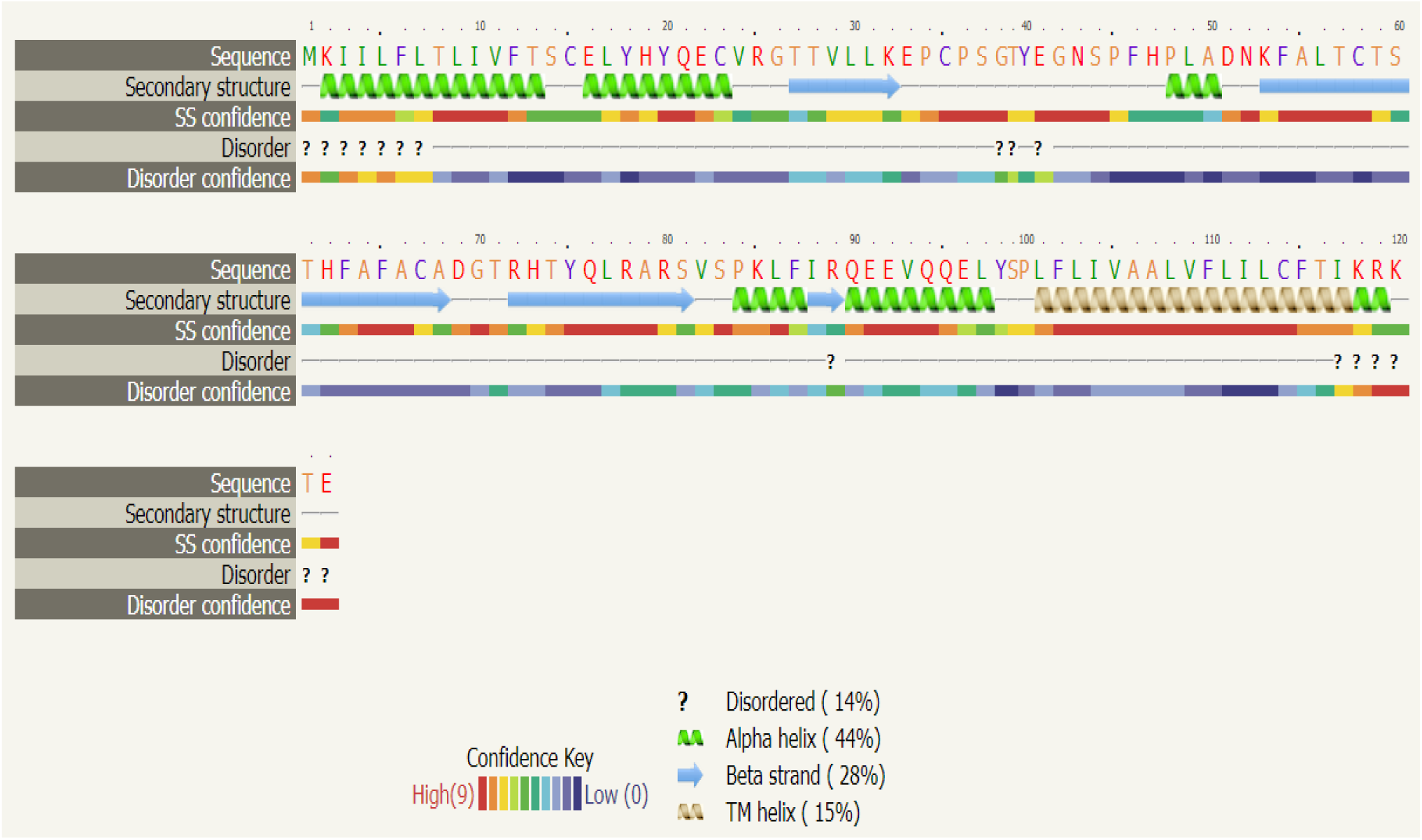
phyre2 results for secondary structure and disorder prediction for ORF7a

The top line displays the place in the order. The sequence is shown on the next line, color-coded residues using a simple property-based system. The three possible states are α-helix, β-strand, or coil. The -helices are represented by green helices, the -strands by blue arrows, and the coil by thin lines.

The “SS confidence” line, with red denoting a high level of confidence and blue denoting a low one, shows the confidence level in the forecast. The forecast of disordered areas in your protein is contained in the “Disorder” line, and these regions are denoted by question marks (?). Disordered areas can frequently be crucial in terms of functionality. However, there is almost no point in forecasting their structure, considering how disordered they appear. ORF7a is indicated here with 14% disorder, 44% helix, 28% beta-strand, and 15% Transmembrane helix (TM helix).

To evaluate if your sequence is likely to include transmembrane helices and forecast their topology in the membrane, a Support Vector Machine (a potent machine learning tool) is used to analyze your sequence and the collection of homologs discovered by PSI-Blast. Memsat-svm, which Phyre2 employs for this purpose and has an average accuracy of 89 % on a large test set, is used. ORF7a, a transmembrane protein of the SARS-corona virus, has a TM helix and is predicted to have the following structure, as is well documented.

**Figure 11:**
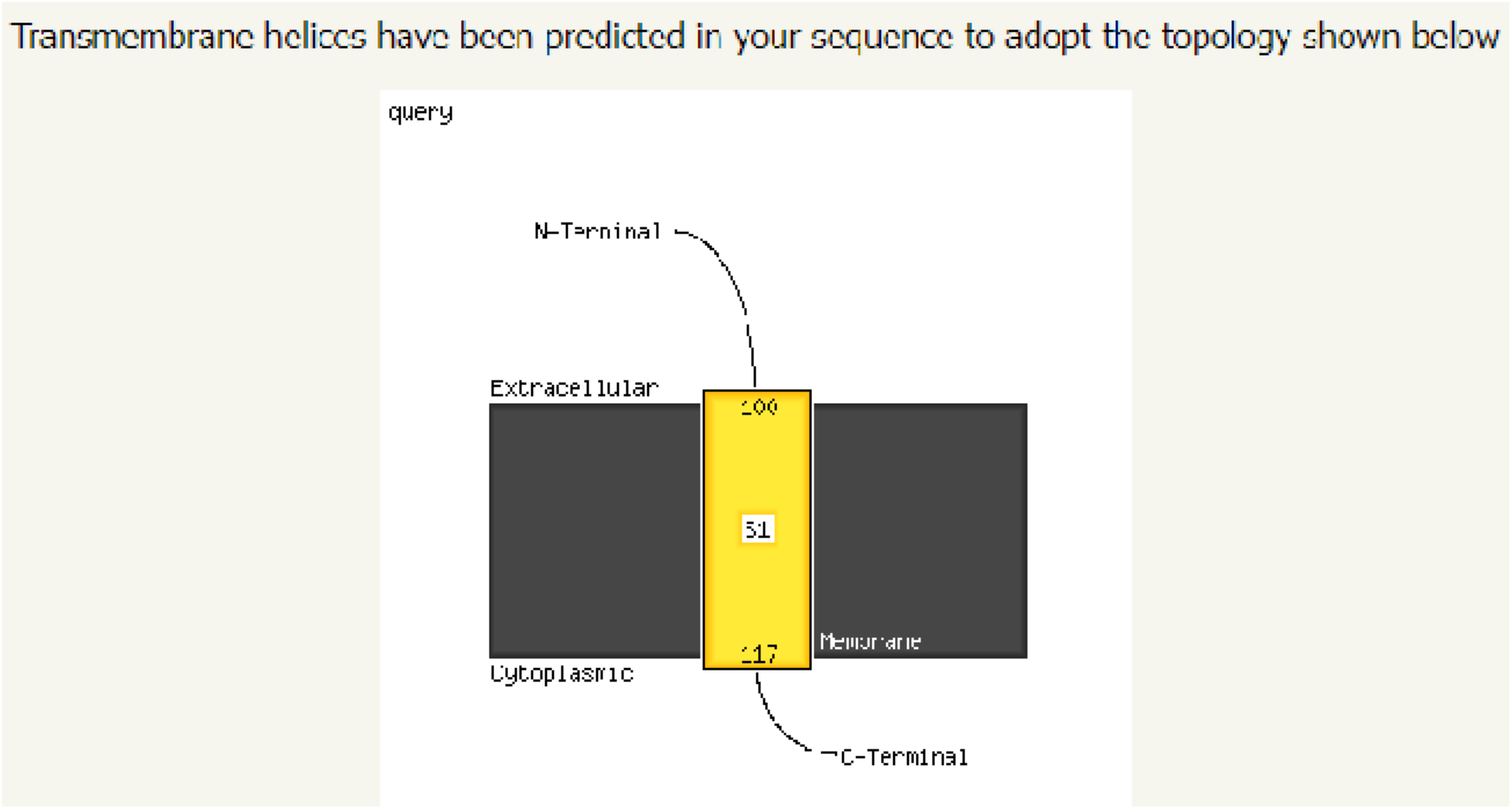
Transmemebrane helix region protein of SARS-CoV-2 (ORF7a) prediction through phyre2

The confidence level of the color coding used in the domain analysis section shows where your sequence matches have been discovered. The red line denotes a strong (red) match to the protein fold library over the whole protein length. The letter “c” denotes that this protein is a complete chain retrieved from the PDB. The domain is denoted by the letter “d.” Below these hits, the colors shift to green and blue, signifying a decline in match confidence.

**Figure 12:**
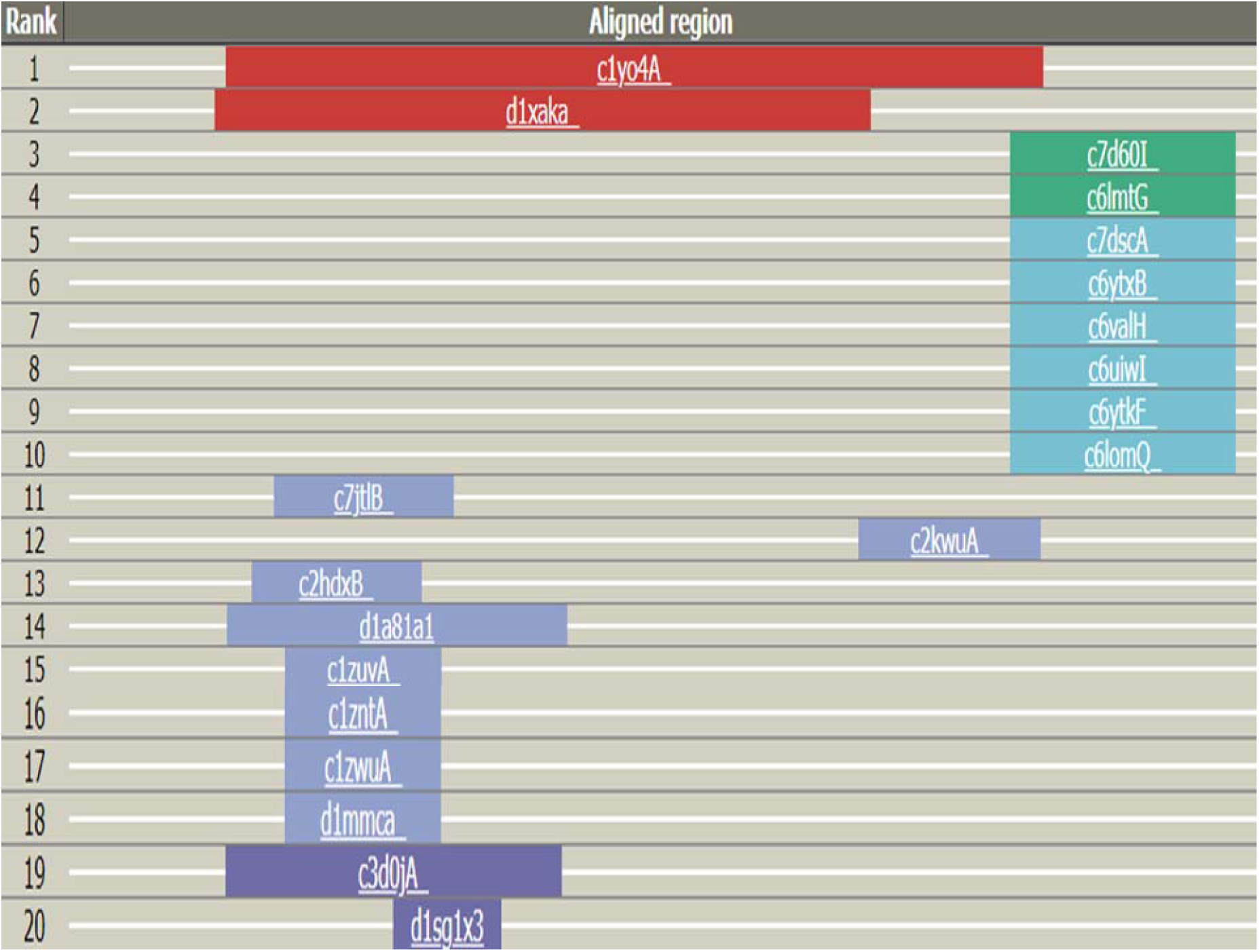
domain alignment through phyre2 of SARS-CoV-2 protein (ORF7a)

A raw alignment score that considers the quantity and quality of aligned residues is used to rank the matches. The similarity of the residue probability distributions at each position, the similarity of the secondary structures, and the presence or absence of insertions and deletions all contribute to this. Two models with templates c1yo4A and d1xaka, with a confidence level of 100 %, are the top two models based on alignment for ORF7a.

**Figure 13:**
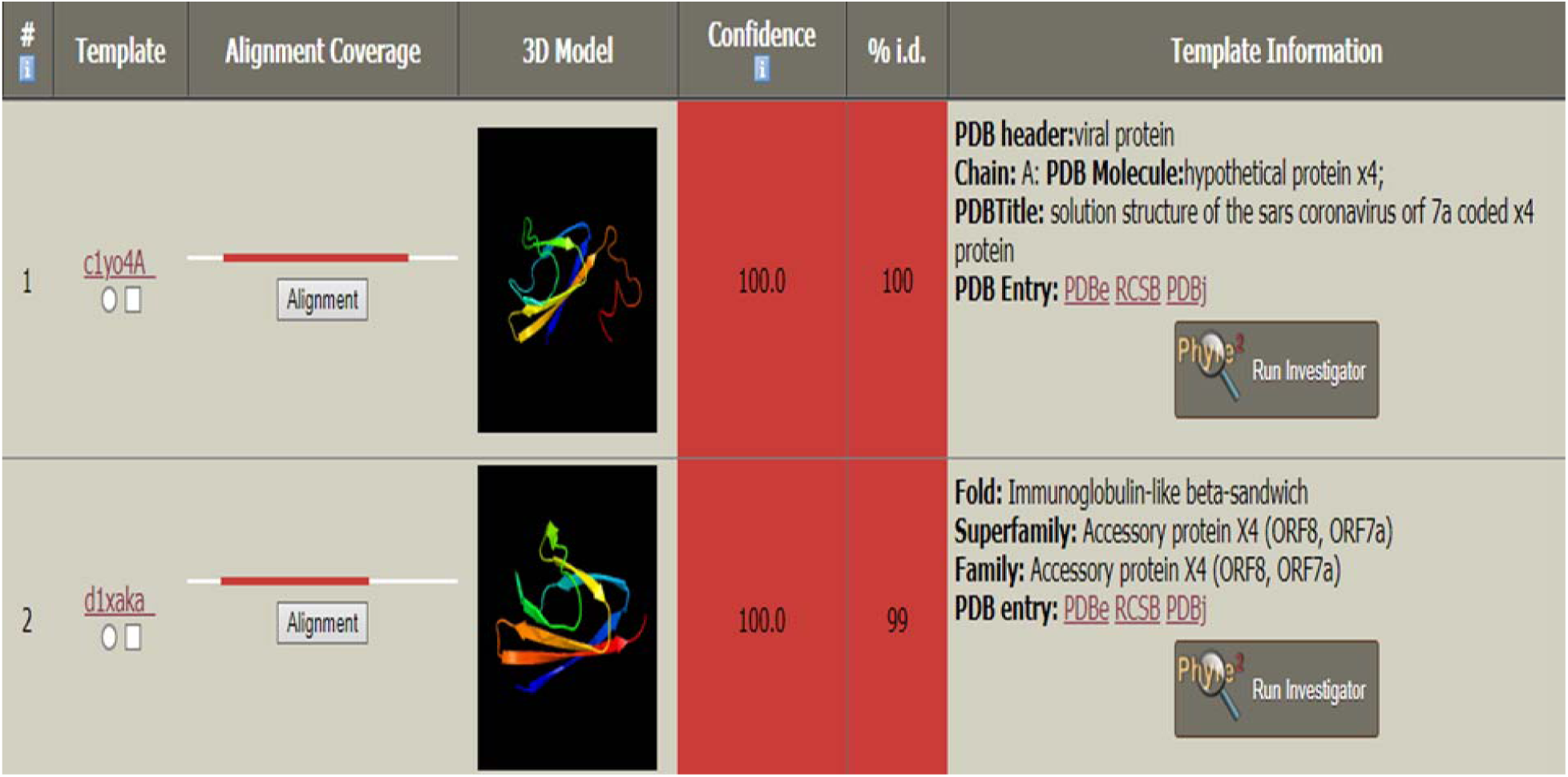
detailed alignment of top 2 templates which closely relates with ORF7a, protein of SARS-CoV-2

### 4.2. IntFOLD

The IntFOLD service offers a single point of access for the automated prediction of tertiary protein structures with built-in estimations of model accuracy, protein structural domain borders, naturally disordered protein regions, and protein-ligand interactions (McGuffin et al., 2019). Using a single-template local consensus fold recognition methodology, the IntFOLD-TS technique predicted the protein tertiary structure from sequence. Since then, the technique has been upgraded to use a cutting-edge multiple-template modeling strategy influenced by both global and local quality estimations. The most recent version includes more sequence-structure alignment techniques, which increases model accuracy even more (Debnath et al., 2022).

Based on independent official assessment measures, the IntFOLD server has established itself as one of the top-performing publically accessible servers. An integrated collection of high-performance, completely automated tools for predicting the structure and function of proteins from their amino acid sequences are freely accessible through the IntFOLD website. The Team of CAMEO (Continuous Automated Model EvaluatOn) regularly benchmarked the server’s component methods. Recent CASP studies have thoroughly blind-tested them. The IntFOLD algorithms have been independently proven to be among the best servers in several prediction categories (McGuffin et al., 2021, McGuffin et al., 2019).

With graphical output offering a visual summary of a complicated data set, the IntFOLD server’s results are understandably given to non-expert users. Individual forecasts more specific outcomes may be interactively inspected, and standard data formats can be used to obtain the unprocessed, machine-readable data (McGuffin et al., 2019).

#### 4.1.3 Methodology

Like other servers, IntFOLD requires only the protein amino acid sequence for its tertiary structure prediction. The sequence of ORF7a is inserted in the search bar of the website. After 24 hours, the result will be displayed on the website, and an email will be sent when the prediction is completed.

#### 4.1.4 Results

IntFOLD creates 5 top models along with their local model quality plot, global quality model score, confidence, and P-value. Results are ordered in decreasing order of overall model quality. The rating for the overall model quality falls between 0 and 1. Scores of less than 0.2 often suggest that certain domains may have been modeled inaccurately. In contrast, scores of more than 0.4 typically denote more thorough and confident models comparable to the native structure.

The global scores consistency enables us to determine a p-value, which is the likelihood that any model is flawed. In other words, the p-value shows the percentage of wrong models with a certain projected model quality score. Depending on the p-value, each model is also given a confidence level with a color code.

**Table 4:** Confidence and P-value

| P-value cut-off | Confidence | Description |
| --- | --- | --- |
| $p < 0.001$ | **CERT** | Less than 1/1000 incorrect models will have higher scores. |
| $p < 0.01$ | **HIGH** | Less than 1/100 incorrect models will have higher scores. |
| $p < 0.05$ | **MEDIUM** | Less than 1/20 incorrect models will have higher scores. |
| $p < 0.1$ | **LOW** | Less than a 1/10 incorrect models will have higher scores. |
| $p > 0.1$ | **UNCERT** | More than 1/10 incorrect models will have higher scores. |

The confidence scores should be considered along with the quality of the local model and the coverage of the target protein by the template or templates. Each row in the results table contains thumbnail pictures of charts showing the per-residue error versus residue number. IntFOLD designed several models, but only the top 5 were selected through quality assessment. Additionally, a thumbnail of the model’s 3D perspective is shown in each table row. Furthermore, the table along with the model ID is mentioned.

**Table 5:**
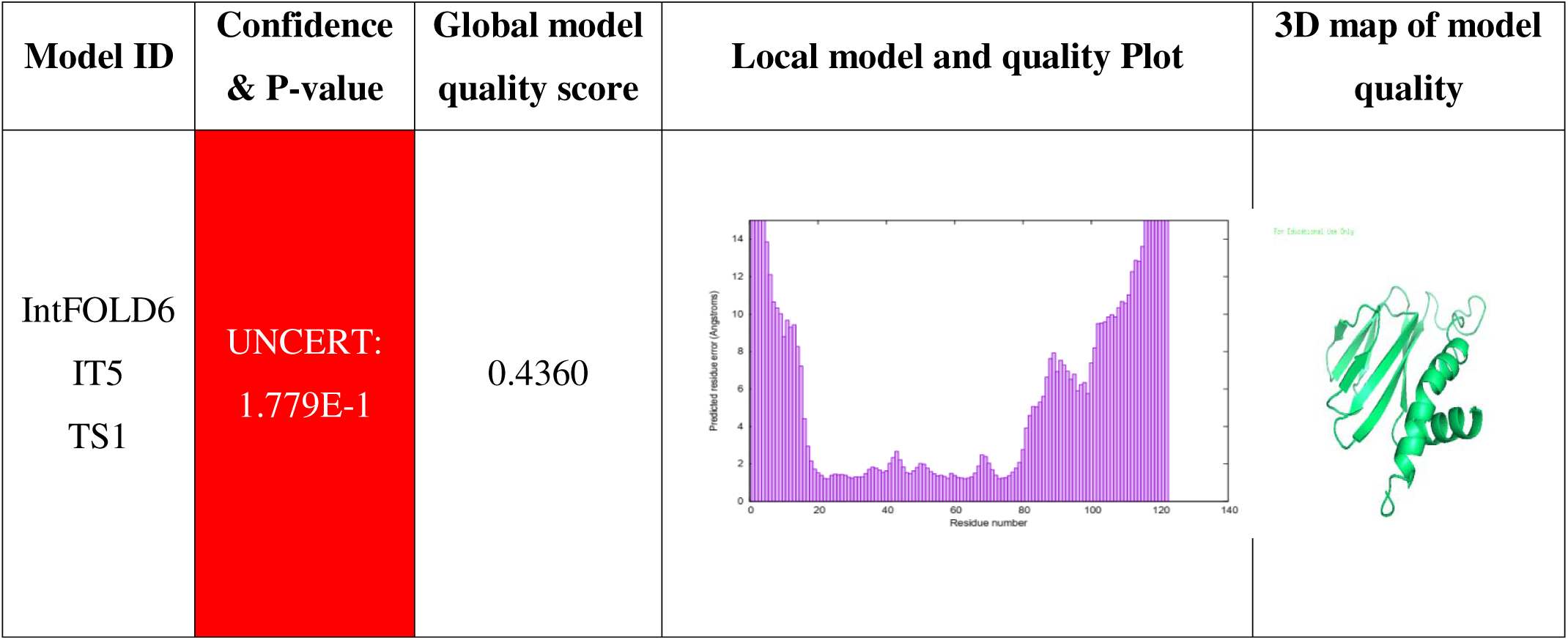
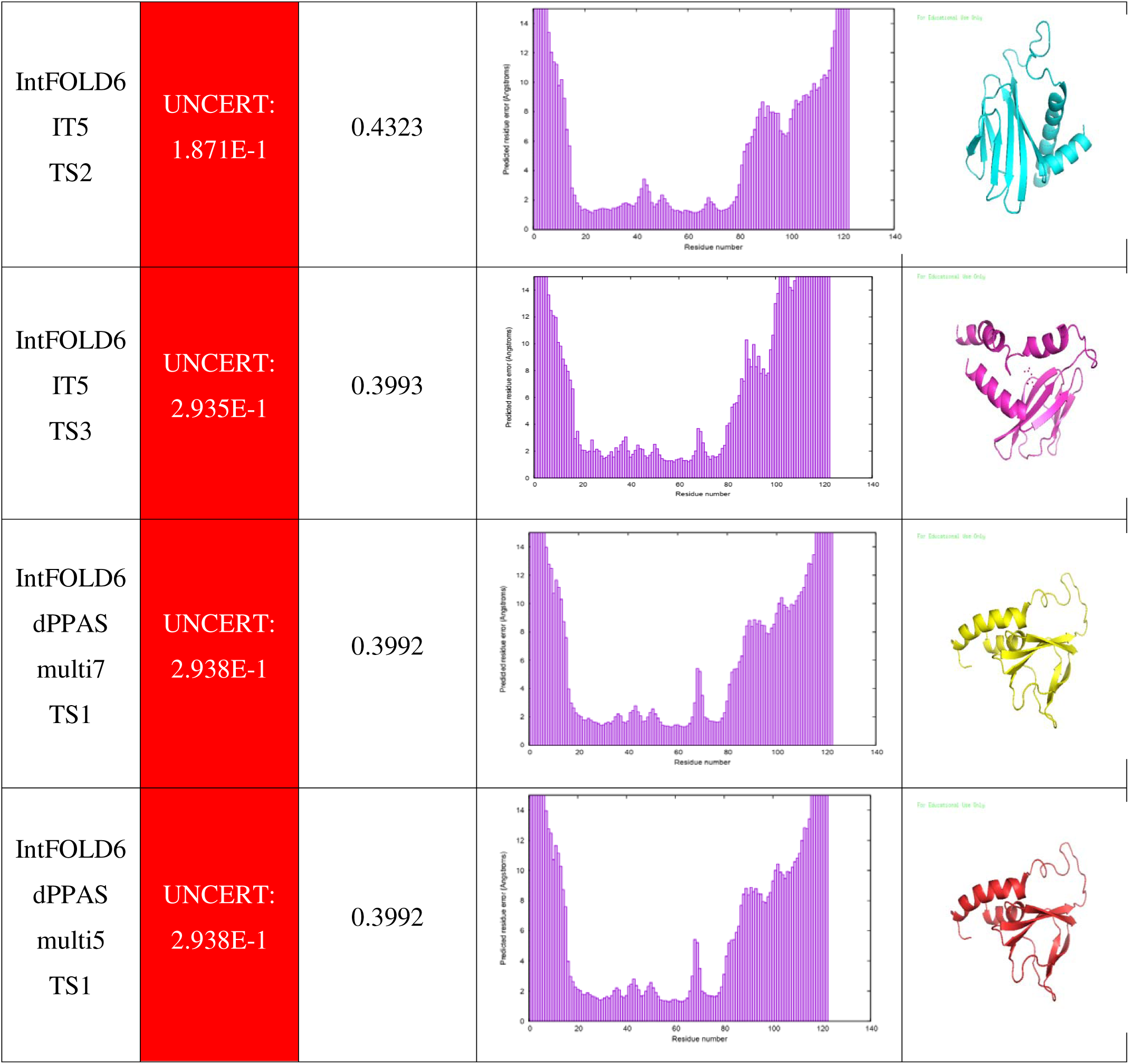
Tertiary structure prediction of top 5 models for ORF7a

| Model ID | Confidence & P-value | Global model quality score | Local model and quality Plot | 3D map of model quality |
| --- | --- | --- | --- | --- |
| IntFOLD6<br>IT5<br>TS1 | **UNCERT:**<br>**1.779E-1** | 0.4360 |  |  |
| IntFOLD6<br>IT5<br>TS2 | UNCERT:<br>1.871E-1 | 0.4323 |  |  |
| IntFOLD6<br>IT5<br>TS3 | UNCERT:<br>2.935E-1 | 0.3993 |  |  |
| IntFOLD6<br>dPPAS<br>multi7<br>TS1 | UNCERT:<br>2.938E-1 | 0.3992 |  |  |
| IntFOLD6<br>dPPAS<br>multi5<br>TS1 | UNCERT:<br>2.938E-1 | 0.3992 |  |  |

The graph displays a probability of disorder (on the y-axis) plot for each numbered amino acid in the sequence (on the x-axis). The graphic displays the disorder/order probability threshold as a dashed line. Residues below the threshold might be considered generally orderly, while those above it are mostly disordered.

**Figure 14:**
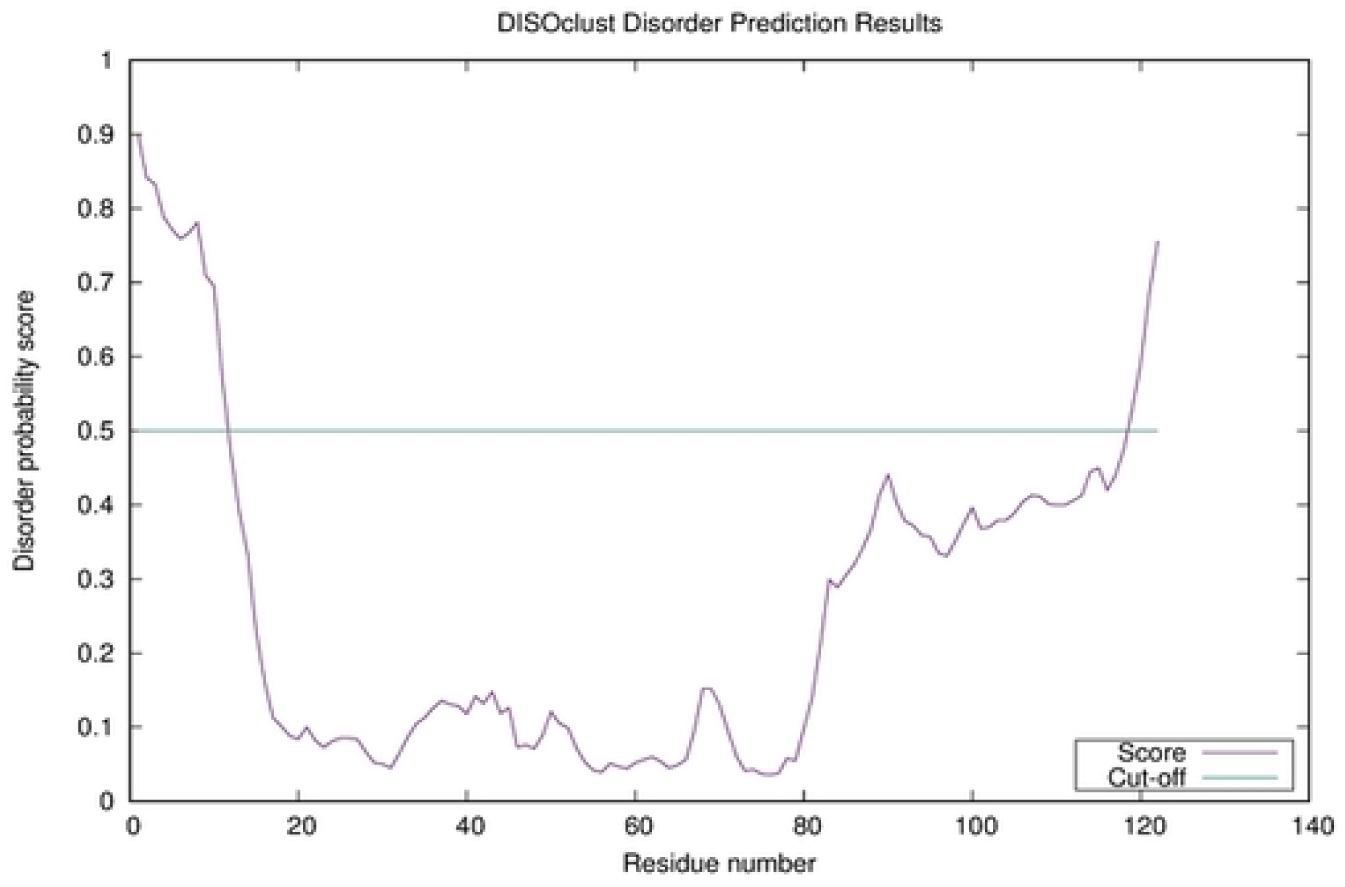
Plot for disorder probability score versus residue number

ORF7a is a transmembrane and has a single domain. As we know, the basic unit of tertiary structure is a domain, which has a unique hydrophobic core made of secondary structural units linked by loop regions. So, the following image shows that the top predicted model of ORF7a consists of only one domain.

**Figure 15:**
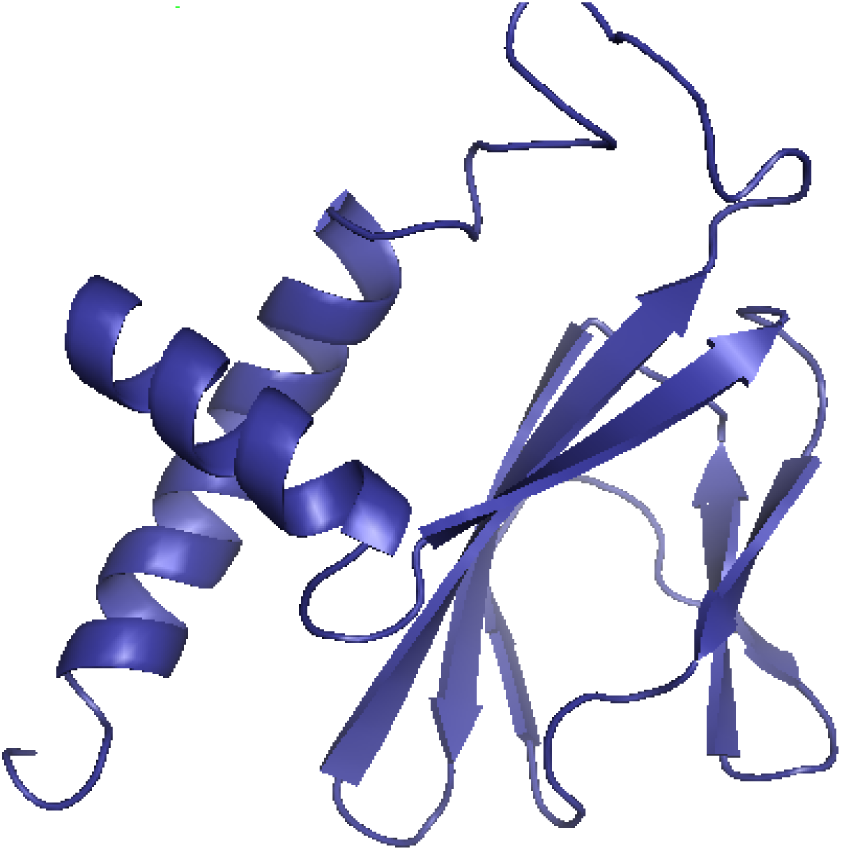
domain prediction for ORF7a top model

A binding site is a location on a macromolecule, such as a protein that specifically attaches to another molecule. A ligand is a common name for the macromolecule’s binding partner. Perhaps, ORF7a lacks a ligand binding site.

### 4.3 Robetta

The University of Washington’s Baker lab created Robetta, a server for predicting protein structures. Comparative modeling (homology modeling) or the de novo fragment insertion strategy used by Rosetta can be used to estimate the structure. Methods based on deep learning, such as RoseTTAFold and TrRosetta, have the advantage of being relatively quick and accurate (Baek et al., 2021).

Automated protein structure analysis and prediction methods are available on the Robetta server. Sequences provided to the server are processed into potential domains for structure prediction, and structural models are created. Using BLAST, PSI-BLAST, FFAS03, or 3D-Jury, if a confident match to a protein with a known structure is discovered, it is utilized as a template for comparative modeling. If no match is discovered, structure predictions are created using the de novo Rosetta fragment insertion approach (Ovchinnikov et al., 2016).

In the presence or lack of sequence homology to proteins with known structures, the server employs the first completely automated structure prediction approach to provide a model for a whole protein sequence. The findings are supplied as domain predictions and molecular coordinates of models covering the query. Robetta also offers the option to use “computational interface alanine scanning” to locate energetically significant side chains involved in the interface of protein-protein complexes. On a single CPU, computational alanine scanning is a rapid process that takes only a few minutes to complete (Du et al., 2021).

Robetta uses a fully automated implementation of the Rosetta software package for protein structure prediction (Du et al., 2021). Roberta’s ultimate objective is to offer quality structural data that will support research, predict function, and help with drug design (Yu et al., 2021).

#### 4.3.1 Methodology

First of all, it is necessary to register before submitting the sequence to Robetta. Access the submit page after creating an account and are logged in. After signing in, one can set custom distance constraints, upload templates and alignments for comparative modeling, paste or upload your protein’s amino acid sequence, and more using an interactive interface on the submit page.

#### 4.3.2 Results

As we know, that robetta offers homology modeling and de novo. However, here we are checking our results of ORF7a for ab inito (de novo) modeling. As Robetta is predicting protein structure by ab initio modeling thus, it uses other web servers to assist in protein prediction without a template. Also, it serves as a transmembrane predictor for our protein. RaptorX Property, a web server, uses no template data to predict the structure of a protein sequence. To anticipa e secondary structure (SS), solvent accessibility (ACC), and disorder areas (DISO), this site uses an emerging machine learning technique known as DeepCNF/deepconcnf (Deep Convolutional Neural Fields).

Numerous protein structure prediction techniques are available through the PSIPRED Workbench. It helps with domain and function prediction and secondary structure prediction, including regions of disorder and transmembrane helix packing. SPIDER3 improves the prediction of protein secondary structure, backbone angles, contact numbers, and solvent accessibility by capturing non-local interactions through long short-term memory bidirectional recurrent neural networks. We can also identify disordered regions in protein structures using the Psipred workbench (disopred). TMHMM is a transmembrane predictor.

**Figure 16:**
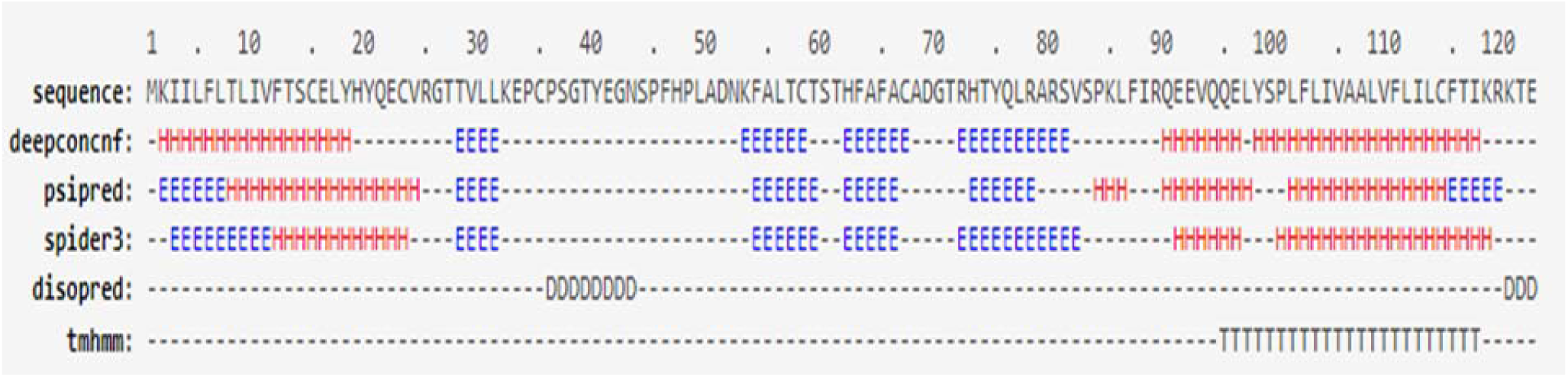
secondary structure and disorder region prediction of ORF7a through Robetta

Through ab initio modeling, Robetta provided us with five top predicted models of ORF7a with estimated angstroms error and the positions of amino acids in the model. De novo models are built using the Rosetta denovo protocol while the procedure is fully automated. Five models, along with their plotted graphs, are mentioned.

**Figure 17:**
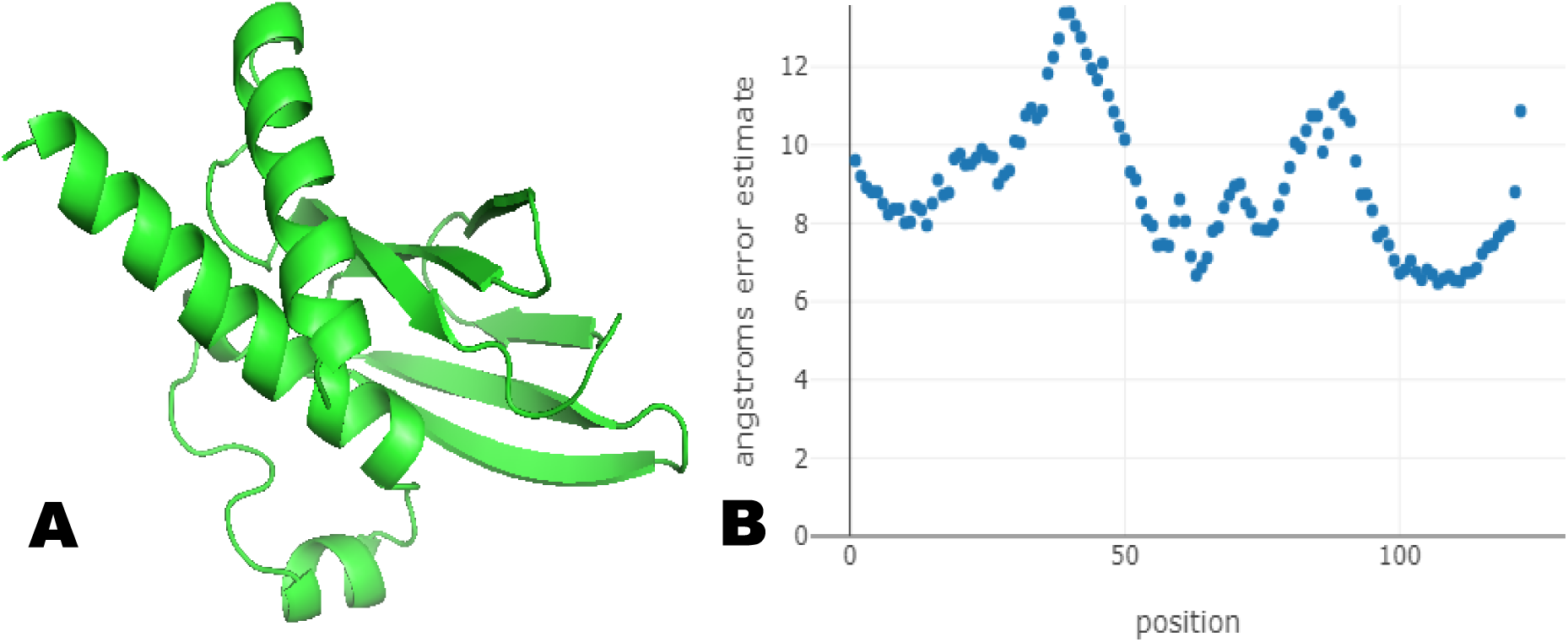
Model 1 with estimated error graph

**Figure 18:**
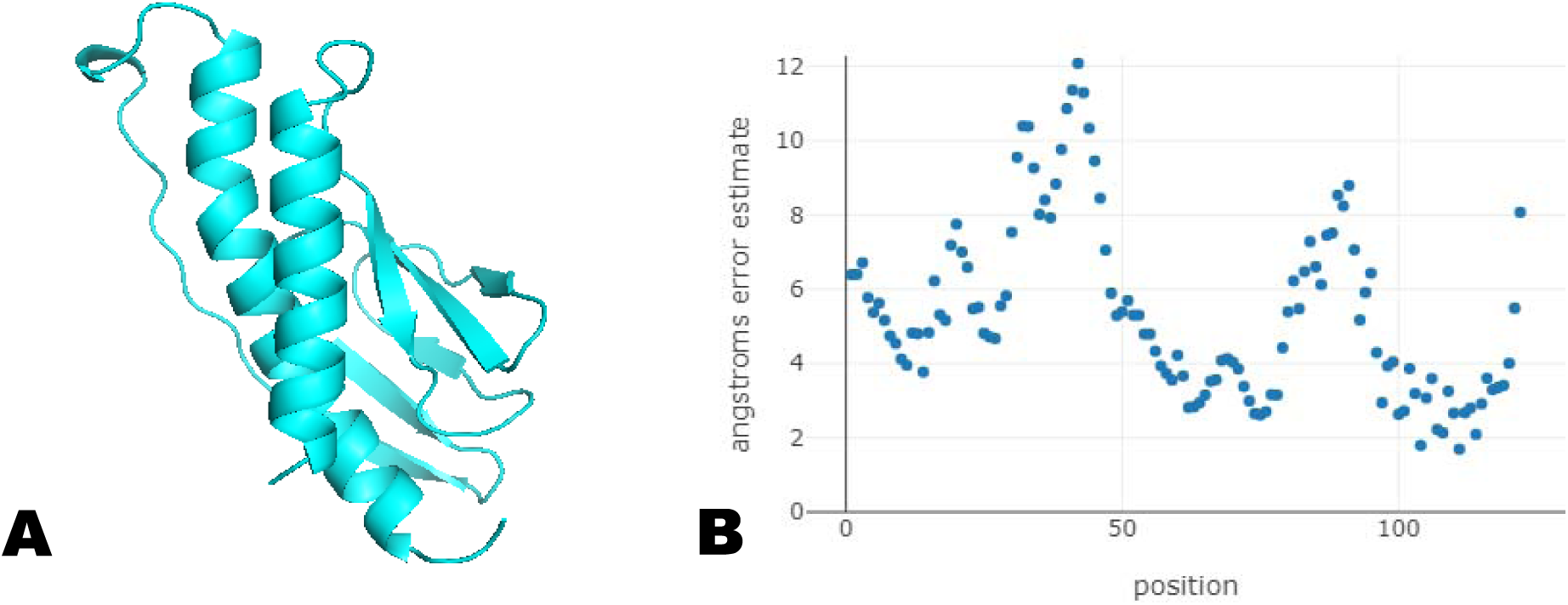
Model 2 along with the plotted graph of the estimated error

**Figure 19:**
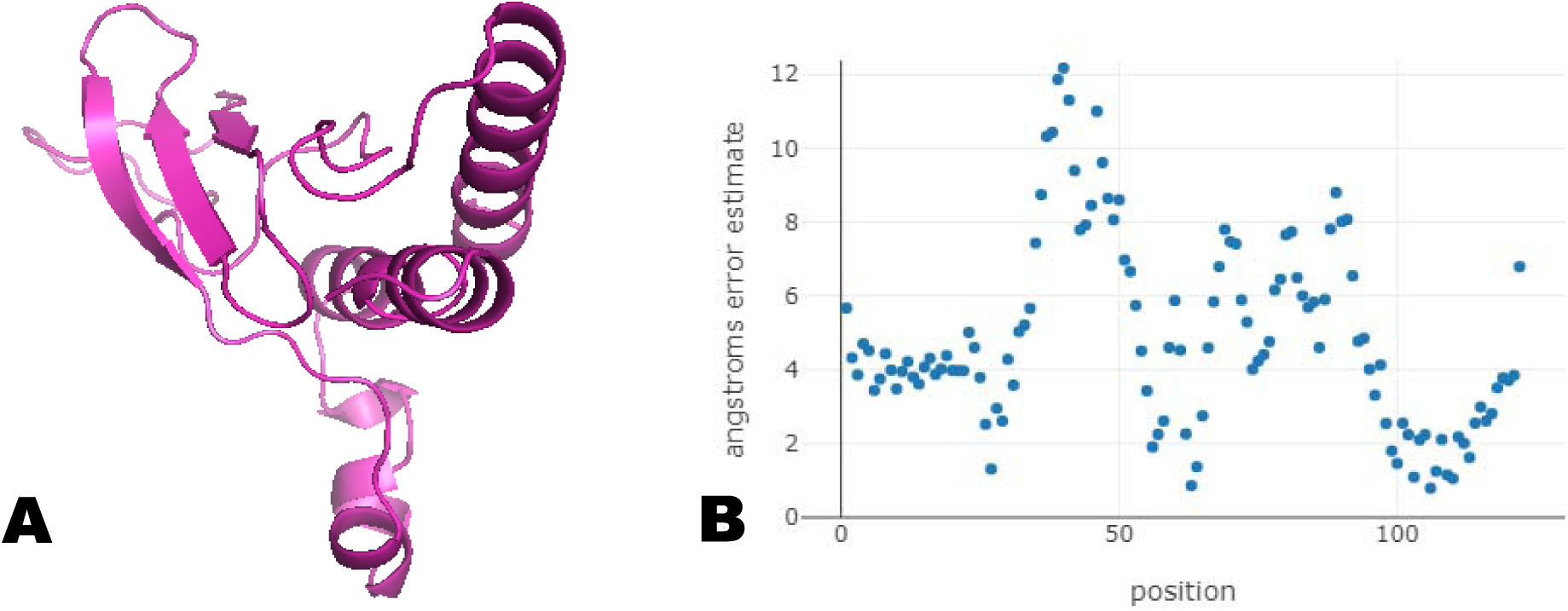
Model 3 with error graph

**Figure 20:**
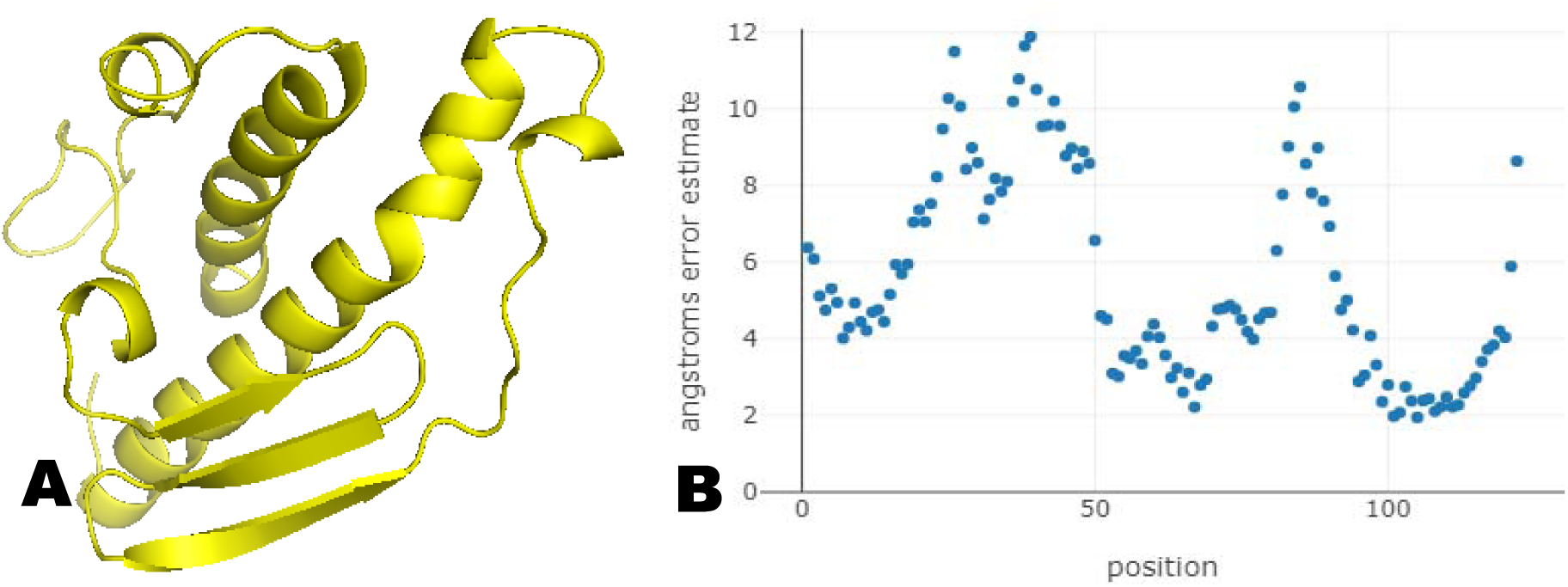
Model 4 with angstroms estimated error graph

**Figure 21:**
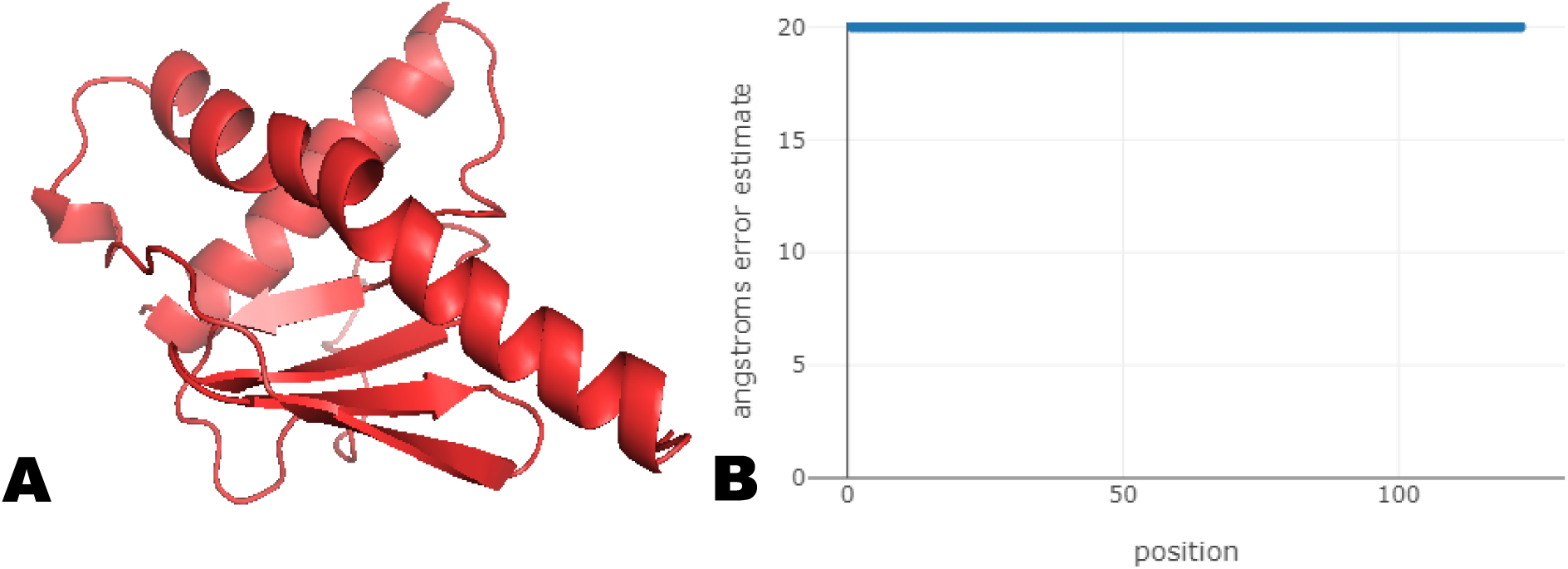
Model 5 with the graph for estimated error

Domain prediction for ORF7a shows that there is one domain in protein. Detecting signals of protein structural domains from sequencing data is known as domain parsing, which also gives results for one domain. The confidence ratio for domain prediction is 0.24. The average TM-score (Template modeling score) of the top 10 Rosetta scoring models reflects the confidence for ab initio domains. It was discovered that these parameters correlated with the real GDT (Global distance test) of the native structure. The confidence matches DeepAccNet-predicted IDDT (local distance difference test) for the TrRosetta and RoseTTAFold domains.

### 5.0 Current challenges and future directions in Protein Structure prediction

The atoms’ configuration in a protein’s three-dimensional structure determines its biological activity. This may refer to how a protein interacts with other proteins for structural or other regulatory functions, or it may refer to the configuration of catalytic residues in an active site. With a better understanding of protein structure and function, we may develop theories about how to influence, regulate, or change it. For instance, elaborating on a protein’s structure might help in protein engineering and site-directed mutation design. A protein’s binding molecules might also be predicted. We can learn about host-antigen interactions, the pathogenesis of any disease, and how to treat it (Marx, 2022, Li et al., 2013).

The protein structure prediction challenge is determined by the evolutionary or structural distance between the target and the solved proteins in the PDB collection because a full physicochemical description of protein folding principles does not yet exist. Full-length models of proteins with close templates can be created by reproducing the template framework. Free modeling is the ‘Holy Grail’ of protein structure prediction since its achievement would signal the problem’s ultimate resolution. Although a solely physics-based Ab initio simulation has the benefit of illuminating the protein folding mechanism, the best current free-modeling findings come from systems that mix both knowledge- and physics-based approaches. Although proper topology for small proteins is consistently achieved, the fascinating high-resolution free modeling is rare and computationally costly (Koehler Leman et al., 2015, Lee et al., 2017).

Nonetheless, there are many problems and challenges to overcome, such as (i) the performance of current alignment algorithms for remote homology templates needs to be improved; (ii) the force field for conformational search is not accurate enough or is not optimal for the specific protein representation; (iii) the global topology and local structure are difficult to refine simultaneously; membrane proteins, whose structures are difficult to identify by experiment and are equally challenging to forecast computationally; (v) the prediction of protein complexes (quaternary structure) is crucial but currently lacking. We think that all of these issues will be overcome as the study progresses. Furthermore, as the CASP experiment organizers pointed out, most existing prediction systems depend too much on known structural information, which is exactly what we do not want to observe. A breakthrough in protein structure prediction is anticipated by minimizing reliance on existing structures and improving first-principles research. We anticipate that the second genetic code will be deciphered by protein structure prediction shortly (Deng et al., 2018, Dunkelmann et al., 2021).

## Conclusion

Protein structure prediction is now one of the most representative and important research fields in computational biology and bioinformatics. The current overview briefly introduces protein structure prediction, along with its background, prediction methodology, general procedure, online web servers, and methods for protein structure searches. Many prediction techniques are currently in use by molecular biologists worldwide and are already very developed. We hope that readers will find it useful by providing a general overview of protein structure prediction. Moreover, it is possible to forecast the new development in protein structure prediction by lowering reliance on existing structures and fostering first-principles research.

In addition, we employ bioinformatics structure prediction tools to elucidate the structure and domain of ORF7a, which is named after the gene that encodes it. It is a transmembrane protein of SARS-CoV-2 that triggers a strong immunological response. It interacts with CD14+ monocytes and enhances the production of proinflammatory cytokines while decreasing the level of Human leukocyte antigen expression. We can treat SARS-CoV-2 and discover a means to eradicate it from our environment by understanding the illness’s virulence components and disease mechanism. However, some of the development in protein structure prediction we have discussed here is due to the growth in the number of protein sequences and structures and the accessibility of computing power. Having the right tools when predicting structures is crucial to assessing your confidence level in the resulting models. However, novel and enhanced methods did play a role in accurately forecasting the major advancement structure. It is still hard to forecast the structure of proteins, and more work is still needed to be done.

## Acknowledgment

The authors of the current article are thankful to Dr. Syeda Mahreen-ul-Hassan Kazmi and the team of BigBio for their kind supervision and throughout support.

